# Self-regulation of stress-related large-scale brain network balance using real-time fMRI Neurofeedback

**DOI:** 10.1101/2021.04.12.439440

**Authors:** Florian Krause, Nikos Kogias, Martin Krentz, Michael Lührs, Rainer Goebel, Erno J. Hermans

## Abstract

It has recently been shown that acute stress affects the allocation of neural resources between large-scale brain networks, and the balance between the executive control network and the salience network in particular. Maladaptation of this dynamic resource reallocation process is thought to play a major role in stress-related psychopathology, suggesting that stress resilience may be determined by the retained ability to adaptively reallocate neural resources between these two networks. Actively training this ability could hence be a potentially promising way to increase resilience in individuals at risk for developing stress-related symptomatology. Using real-time functional Magnetic Resonance Imaging, the current study investigated whether individuals can learn to self-regulate stress-related large-scale network balance. Participants were engaged in a bidirectional and implicit real-time fMRI neurofeedback paradigm in which they were intermittently provided with a visual representation of the difference signal between the average activation of the salience and executive control networks, and tasked with attempting to self-regulate this signal. Our results show that, given feedback about their performance over three training sessions, participants were able to (1) learn strategies to differentially control the balance between SN and ECN activation on demand, as well as (2) successfully transfer this newly learned skill to a situation where they (a) did not receive any feedback anymore, and (b) were exposed to an acute stressor in form of the prospect of a mild electric stimulation. The current study hence constitutes an important first successful demonstration of neurofeedback training based on stress-related large-scale network balance – a novel approach that has the potential to train control over the central response to stressors in real-life and could build the foundation for future clinical interventions that aim at increasing resilience.

**Highlights:**

- Acute stress affects the allocation of neural resources between large-scale brain networks
- We provide a first successful demonstration of neurofeedback training based on stress-related large-scale brain networks
- Novel approach has the potential to train control over central response to stressors in real-life
- Could build foundation for future clinical interventions to increase resilience

## 1 Introduction

Our body’s response to acute stress constitutes an essential adaptive mechanism that helps us to properly evaluate and react to potential threats in our environment (de Kloet et al., 2005). However, repeated exposure to stressors can also lead to maladaptation and mental disorders (Kalisch et al., 2015). Stress related mental illness, such as depression and anxiety disorders, are amongst those with the highest disease burden and efforts to reduce their high prevalence have remained largely unsuccessful (Rehm & Shield, 2019; Vos et al., 2017). This has triggered a paradigm shift from treatment to prevention-oriented health research over the last years, with a steadily increasing interest in resilience – an individual’s ability to positively adapt to being exposed to a stressor (Kalisch et al., 2015).

Recent neuroimaging research has revealed how acute stress affects the human brain at the systems level, with stress-related hormones and neurotransmitters triggering shifts in large-scale brain network configurations (Hermans et al., 2014; Hermans et al., 2011; van Oort et al., 2017). In particular, stress appears to induce a shift in the balance between the salience network (SN), which integrates cognitive processes associated with salient stimuli, including bottom-up attention (Seeley et al., 2007), and the executive control network (ECN), which regulates higher-order cognitive functions such as working memory and top-down attention (Vincent et al., 2008). It is believed that during the acute stress phase, neural resources are reallocated to strengthen SN activity at the cost of ECN function (Young et al., 2017) - a balance shift that is subsequently actively reversed to return to homeostasis (Hermans et al., 2014). Maladaptation of this dynamic resource reallocation process is thought to play a major role in stress-related psychopathology (Akiki et al., 2017; van Oort et al., 2017; Menon, 2011), suggesting that stress resilience may be determined by the retained ability to adaptively reallocate neural resources between these two networks. Actively training this ability could hence be a potentially promising way to increase resilience in individuals at risk for developing stress-related symptomatology.

In the current study, we investigated training the voluntary reallocation of neural resources between SN and ECN using real-time fMRI neurofeedback (rtfMRI-NF), as a potential mean to increase stress resilience. Following a short localizer session from which subject-specific network masks were defined, healthy participants were each engaged for three separate sessions in a bidirectional, implicit and intermittent rtfMRI-NF paradigm in which the difference signal between the average activations in the individualized SN and ECN masks was coupled to the size of a visual stimulus on the screen. Participants were not given any details of this setup other than the instruction that they could learn to control the size of the stimulus *with their brain*. In a subsequent transfer session, participants performed the same bidirectional self-regulation task, but had to apply their learned strategies in the absence of any feedback and, in addition, were exposed to an acute stressor (mild electric stimulation) in some of the trials (McMenamin et al, 2014). We hypothesized that participants are able to (1) learn to self-regulate SN-ECN activation balance, and (2) apply learned regulation strategies in (a) the absence of feedback and (b) in prospect of an acute stressor. Through analysis of participants’ self-evaluation as well as additional exploration of whole-brain activations during regulation, we aimed to gain further insights into how the tested group of participants achieved self-regulation of the feedback signal.

## 2 Method

### 2.1 Participants

Eleven healthy volunteers (6 females, 5 males; all recruited at Radboud University and Radboud University Medical Centre, Nijmegen, The Netherlands) aged between 19 and 40 years (mean = 25.73; SD = 5.87) participated in the study in return for a monetary reward of 104 – 129 €. All of them had normal or corrected to normal vision and had no known neurological or psychological disorders. Exclusion criteria included MRI contraindications, such as the presence of electronic or ferromagnetic body implants and a prior history of claustrophobia or panic attacks. Only non-native speakers of English were included, to ensure that all participants were at approximately the same level of language proficiency when receiving instructions and answering questionnaires in English. Before the study, participants received general information about fMRI neurofeedback as well as study-specific information pertaining to the scheduling of the study and a short description of the experimental task, and were informed that the three best performers over the whole study would receive an extra monetary bonus of 25 €. Participants were instructed to refrain from the use of recreational drugs, have a good rest the night before each experimental session, and to abstain from consuming caffeinated or alcoholic drinks, and smoking six hours prior to each experimental session. Due to technical problems, one participant had to be excluded after the second experimental session. The remaining 10 participants (5 females, 5 males; between 19 and 40 years of age, mean = 25.80, SD = 6.18) completed all experimental sessions. The study was approved by the local ethics committee, and participants gave their written informed consent before the procedure.

### 2.2 Design

The study consisted of five consecutive experimental sessions (see Figure 1), each on a different day, with 1 to 19 days between the first and the second session (mean = 7.45, SD = 5.68), 1 to 16 days between the second and the third session (mean = 7.20, SD = 5.67), 1 and 13 days between the third and the fourth session (mean = 5.60, SD = 4.09), and 1 and 7 days between the fourth and the fifth session (mean = 4.30, SD = 2.63). The first session (*Localiser*) entailed an anatomical MRI recording, followed by two functional MRI runs (each 300 volumes), with the first being a resting state run used for individualizing network templates, and the second being a passive viewing of pseudo-randomized presentations of five repetitions (each 10 s) of the experimental stimuli used in later sessions to collect baseline pupil responses. Subsequently, a set of questionnaires was administered outside the MR scanner. The following three sessions (*Training*) were identical to each other, and each comprised an anatomical MRI recording, followed by seven to eight functional MRI neurofeedback runs (each 600 volumes). Each functional run started with a long rest block (34 s), followed by 16 self-regulation blocks (“larger”, “smaller”; each 16 s), each followed by “delay” (6 s), “feedback” (4 s) and “rest” (10 s) blocks. In the first and second run of the first Training session, all self-regulation blocks were of the condition “larger” and “smaller”, respectively, to familiarize participants with the paradigm, while all other runs consisted of an equal number of both conditions in a pseudo-randomized order. The fifth and last session (*Transfer*) involved an anatomical MRI recording, followed by four functional MRI runs, and was included to examine if participants were able to apply learned strategies in the absence of feedback, as well as during the presence of an acute stressor. The first three functional runs entailed self-regulation without feedback (762, 710 and 736 volumes, respectively), with a pseudo-randomized order of “larger”, “smaller”, “larger|threat”, “smaller|threat” and “rest|threat” blocks (each 16 s), intermixed with “rest” blocks (each 10 s). Over all three runs, 10% of the threat blocks (“smaller”|threat”, “larger|threat”, “rest|threat”) were replaced by a shock block (“smaller|shock”, “larger|shock”, “rest|shock”), respectively. The first run included one replacement for each of the three shock condition, equally spaced over the whole run, with the “rest|shock” block in the middle. In the second run there was one replacement of “rest|shock” block at approximately the middle of the run. The last run included one replacement for each “rest|smaller” and “rest|larger” at the first and last quarter of the run, respectively. The order of the two replacements for “rest|smaller” and “rest|larger” in the first and third run was counterbalanced over participants to control for order effects. The fourth and last functional run was a resting-state run (300 volumes).

**Figure 1.**
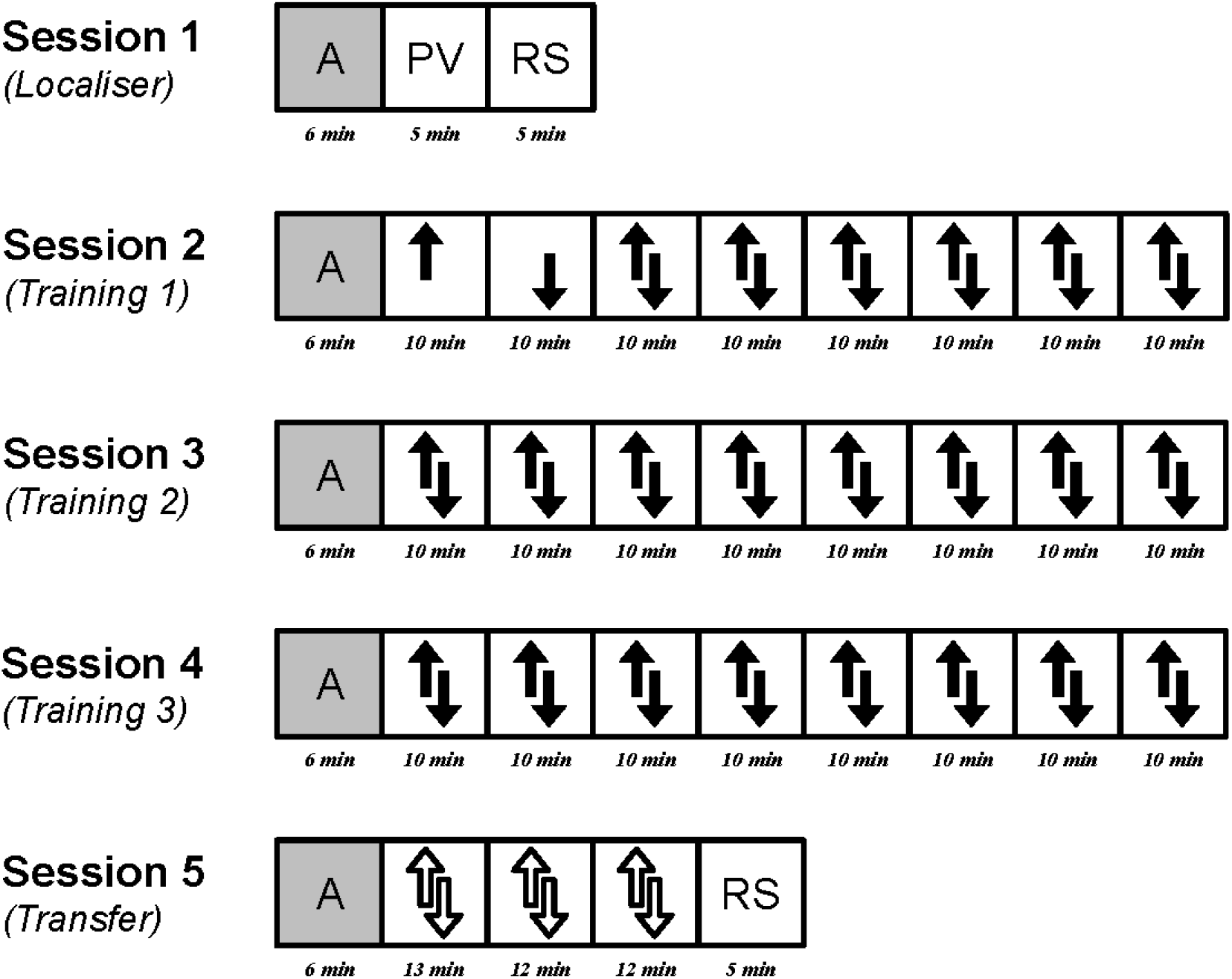
Overview of MRI runs during each of the five experimental sessions. Each session started with the acquisition of an anatomical recording (A). The first session (Localiser) included two short functional runs: passive viewing of the experimental stimuli (PV) and resting state (RS). The second session (Training 1) started with two functional runs in which participants were asked to either increase and decrease the circle, respectively, followed by up to six further runs in which they were asked to do both, in a mixed order, and participants received feedback after each attempt. The next two sessions (Training 2 and Training 3) each consisted of up to eight of these mixed runs. In the last session (Transfer) participants were asked to increase and decrease the size of the circle with the strategies learned during the training, but without receiving feedback and with the prospect of receiving a mild electric stimulation in 50% of the trials. The session ended with a resting state run (not analysed).

### 2.3 Materials

#### 2.3.1 Questionnaires

Administered questionnaires included the 60 item International Personality Item Pool NEO Questionnaire (IPIP-NEO-60; Maples-Keller et al., 2019), the behavioural inhibition system (BIS) and behavioural activation system (BAS) questionnaires (Carver & White, 1994), the cognitive emotional regulation questionnaire (C.E.R.Q.; Garnefski, Kraaij, & Spinhoven, 2001), the trait component of the state-trait anxiety inventory (STAI-trait) (Spielberger, Gorsuch, & Lushene, 1968), the thought control questionnaire (TCQ; Wells & Davies, 1994), and Beck’s depression inventory II (BDI-II; Beck, Steer, & Brown, 1996). Questionnaires were part of a standard battery to identify severely depressed participants (BDI-II), and to add to a larger body of explorative data for generating potential hypotheses for future studies (remaining questionnaires).^1^

#### 2.3.2 Stimuli

Experimental stimuli were created and presented using Expyriment (version 0.9.0; Krause & Lindemann, 2014), running on a computer designated for stimulus presentation. The neurofeedback display consisted of a grey disc (red = 128, green = 128, blue = 128), superimposed with a black circle (red = 0, green = 0, blue = 0; visual angle of radius = 5.96 °, visual angle of thickness = 0.07 °) as well as a black dot in the center (red = 0, green = 0, blue = 0; visual angle of radius = 0.16 °). The size of the grey disc was half between the size of the dot and the size of the black circle (visual angle of radius = 3.06 °) during all blocks except the feedback blocks, during which it could be anything in between size of the black circle and the size of the dot in the center (which turned green in this condition; red =0, green = 255, blue = 0). During regulation blocks, the grey disc was surrounded by four outward (“larger”) or inward (“smaller”) pointing arrows at the top, bottom, left and right side (visual angle of width = 0.77 °, visual angle of height = 0.98 °; visual angle of distances between top and bottom as well as left and right centers = 6.99 °) and the dot in the center turned orange (red = 255, green = 128, blue = 0). During threat blocks, the entire display was additionally surrounded by a red frame (red = 255, green = 0, blue = 0; visual angle of height = 19.36 °, visual angle of width = 19.36 °, visual angle of thickness = 0.33 °). See Figure 2 for an overview of used stimuli.

**Figure 2.**
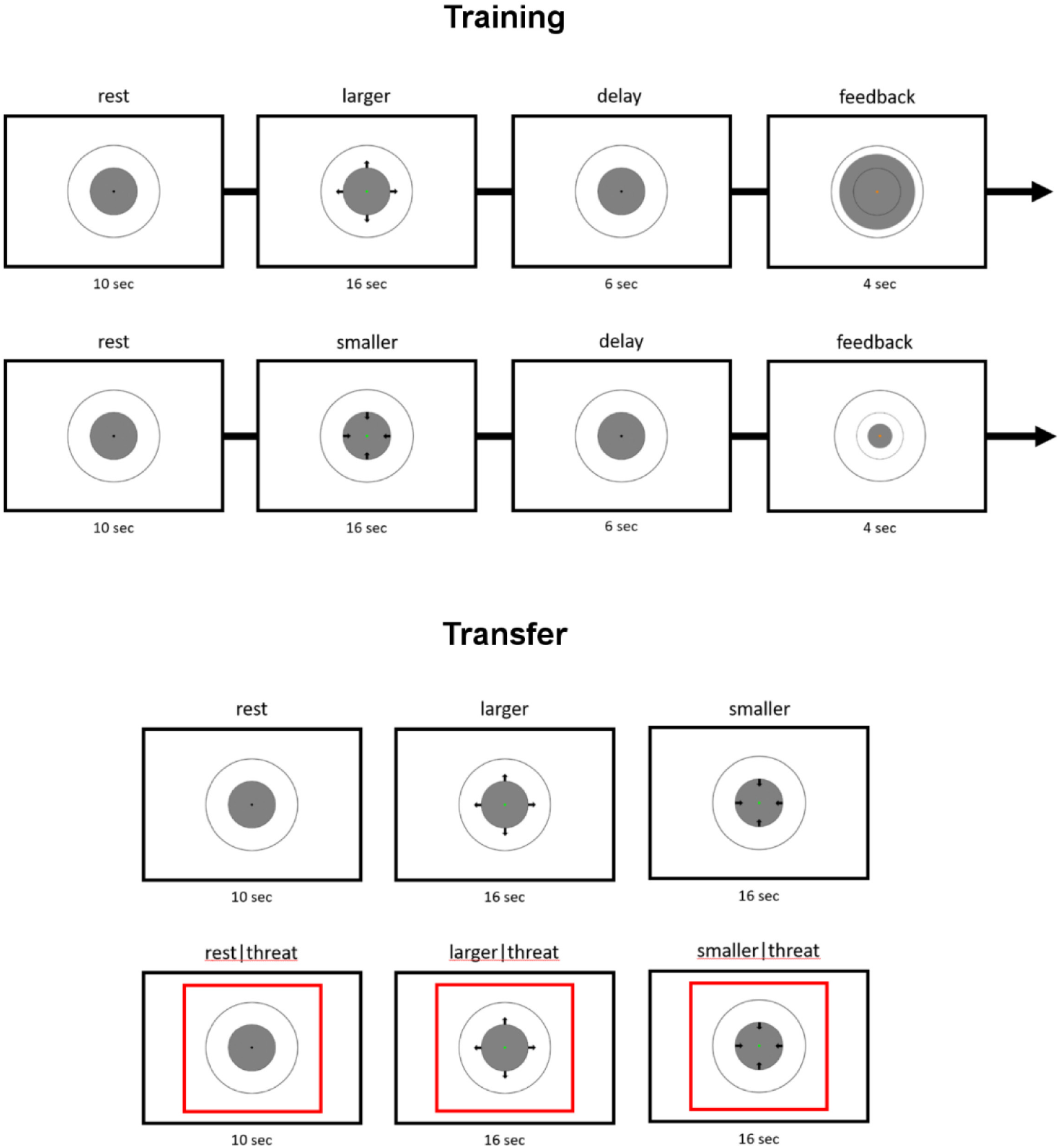
Overview of experimental stimuli and design of the training (top) and transfer (bottom) sessions.

#### 2.3.3 MRI data acquisition

All MR images were recorded using a Siemens Skyra 3T MR system (Siemens, Erlangen, Germany), with a 32-channel receiver head coil. High-resolution 3D anatomical images were recorded using a T1-weighted magnetization-prepared rapid gradient echo (MPRAGE) sequence with a generalized autocalibrating partial parallel acquisition (GRAPPA) acceleration factor of 2 (repetition time/echo time = 2300/3.03 ms, flip angle = 8 °, field of view = 256 × 256 x 192 mm, resolution = 1.0 mm^2^). Functional images were acquired using a echo planar T2*-weighted sequence sensitive to BOLD contrast with a multiband acceleration factor of 4 (repetition time/echo time = 1000/33 ms, flip angle = 60 °, field of view = 210 × 210 mm^2^, number of slices = 52, slice thickness = 2.4 mm (no gap), in-plane resolution = 2.4 x 2.4 mm^2^). During all scans, besides cushioning around the head, a strip of medical tape was applied over the forehead of the participants to reduce head motion (Krause et al., 2019).

#### 2.3.4 Real-time fMRI neurofeedback setup

Real-time functional imaging was realized by implementing a custom functor in the MR image reconstruction pipeline which exported pixel data to an additional computer as soon as it becomes available. TurboExport (version 0.261, Brain Innovation, Maastricht, The Netherlands) was used to transform incoming pixel data for each volume into an image. Each resulting image was preprocessed in real time using Turbo-BrainVoyager (version 4.0 beta; Brain Innovation, Maastricht, The Netherlands). Preprocessing included motion correction (by realigning each image to the first image of the session), as well as spatial smoothing (Gaussian kernel of 5 mm full width at half maximum). The stimulation computer communicated with Turbo-BrainVoyager via a network connection, using the Transmission Control Protocol (TCP), in order to request the preprocessed real-time data to generate the feedback display.

#### 2.3.5 Peripheral recordings

Eye movements and pupil size of the left eye were recorded using an Eyelink-1000 Plus eye-tracker (SR Research, Ottawa, Canada), with a sampling rate of 500 Hz. Additionally, a BrainAmp ExG MR (Brain Products, Gliching, Germany) was used to measure heart rate with an MR-compatible pulse sensor (placed on left ring finger; Brain Products, Gliching, Germany), respiration with an abdominal respiration belt (attached to a pneumatic sensor, Brain Products; Gliching, Germany), and Galvanic Skin Response with two Ag/AgCl electrodes (placed on the distal phalanges of the left index and middle fingers; Brain Products, Gliching, Germany).

#### 2.3.6 Peripheral stimulation

Mild electrical shocks were delivered via two electrodes attached to the first and fifth finger of the right hand using a MAXTENS 2000 TENS unit (Bio-Protech, Gangwon-do, Korea). Stimulation intensity varied between 0 V/0 mA and 40 V/80 mA. During a standardized adjustment procedure prior to the testing, each participant received and subjectively rated five shocks, allowing stimulation strength to converge to an individualized level that was experienced as uncomfortable, but not painful.

### 2.4 Procedure

#### 2.4.1 Session 1: Localiser

After the recording of an anatomical image (≈ 5 min), participants were asked to look at a fixation cross at the centre of the screen, while a resting-state functional data-set was recorded (≈ 6 min). Subsequently, the eye-tracker was calibrated and another short functional data-set was recorded while participants were asked to passively look at the stimuli that were used in the later parts of the study (≈ 5 min). For the purpose of having a baseline pupil size for each of the stimuli, pupil size was recorded during this run. Once the tasks in the scanner were concluded, participants were asked to complete a set of questionnaires (see Materials).

The anatomical image and resting-state data-set were used after the session to create individualized network masks to be used as neurofeedback target regions during the subsequent training and transfer sessions. Using a custom-made Nipype (version 1.1.8; Gorgolewski et al., 2011, Gorgolewski et al., 2017) pipeline (https://github.com/can-lab/IndNet), functional images were realigned to the first volume of the run, spatially smoothed (Gaussian kernel of 5 mm full width at half maximum), cleaned from head-motion artefacts using ICA-AROMA (Pruim et al., 2015), and high-pass filtered (filter size = 100 s). The anatomical image underwent brain extraction and segmentation into grey matter (GM), white matter (WM) and cerebro-spinal fluid (CSF) binary masks. Binary masks of 14 intrinsic connectivity networks (ICN; Shierer et al., 2012) were transformed into native space, multiplied with the GM mask, and the results were used to extract 14 timecourses (first eigenvariate of each ICN) from the cleaned data. In addition, the WM and CSF masks were used to extract average (mean) WM and CSF timecourses from the cleaned data. ICN, WM and CSF timecourses entered a generalized linear model (GLM) in which contrasts reflecting combinations of ICN regressors specifying SN (anterior and posterior), ECN (left and right) as well as the default mode network (DMN; dorsal and ventral) were estimated and thresholded using spatial mixture modelling (threshold level = 0.66). Eventually, SN, ECN and DMN binary masks were created by only considering voxels that occurred exclusively in either of the thresholded maps and transformation into native anatomical voxel space. Resulting masks were imported into BrainVoyager (version 21.0, Brain Innovation, Maastricht, The Netherlands), transformed to be iso-voxeled and a BrainVoyager voxels-of-interest (VOI) definition was created for each mask.

#### 2.4.2 Sessions 2, 3 and 4: Training

At the beginning of the first Training session oral instructions about the task and outline of the session were given outside of the scanner. For the regulation task, participants were asked to attempt to either increase or decrease the size of the disc on the screen with their brain, depending on the orientation of the surrounding arrows in each trial. They were told that they could achieve this by thinking of something specific, performing some mental task internally, or getting into a certain mood, emotion, feeling, or state of mind, and that they had to explore different mental strategies to find one that works for them. They were not made aware of either the origin and computation of the feedback signal, or the details of the study. Before each session, participants were explicitly asked to try to avoid movement, including facial movements, limp movements and irregular breathing patterns. Inside the scanner, first an anatomical image was recorded (≈ 5 min) and preprocessed immediately after reconstruction on a separate computer using BrainVoyager (version 21.0, Brain Innovation, Maastricht, The Netherlands). Preprocessing included intensity inhomogeneity correction, iso-voxeling, and brain extraction. Preprocessed anatomical images were further coregistered to the anatomical image of the first (localiser) session, and results were supplied to Tubro-BrainVoyager to have real-time functional data in alignment with the neurofeedback target ROIs. Parallel to the anatomical preprocessing, the eye tracker was calibrated. Subsequently, participants were engaged in seven (session 4 of participant 3, session 3 of participant 6, session 4 of participant 7) to eight (all other sessions) real-time fMRI neurofeedback training runs (each ≈ 10 min). The main motivation for splitting the sessions into multiple short runs was to offer participants self-paced rest periods (in between runs). Each run started with a rest block during which the baseline and initial display boundaries for the feedback signal (i.e. smallest and largest disc size) were calculated. Feedback was based on the difference signal between the averages (mean) of all voxels in the SN and ECN ROIs (participants 1, 3, 5, 7, 9, 11: SN - ECN; participants 2, 4, 6, 8, 10: ECN - SN). The baseline for this difference signal was defined as the average (median) difference between SN and ECN during the initial rest block, and the initial lower and upper display boundaries were set to two standard deviations from this baseline. These limits were updated before each regulation block to the average (median) of the five lowest/highest difference values in that run. For each regulation block, positive and negative changes in the difference signal resulted in a feedback value between −1 and 1 by applying the following calculation (values lower than −1 and higher than 1 were set to −1 and 1, respectively):

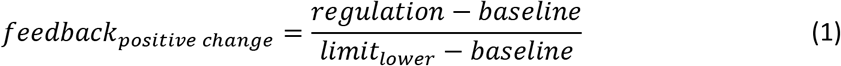

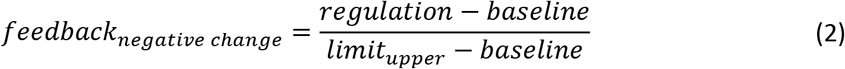

Due to the characteristics of the delayed haemodynamic response, calculations did not consider the first six volumes of each block, but included the first two volumes of the following block. The feedback hence represented the average regulation performance during each regulation block and was given intermittently during the subsequent feedback block. The feedback value was furthermore used to calculate the amount of points a participant collected. For each regulation block, an amount of points proportional to the feedback value was given ranging from 0 points, when the difference signal changed in the opposite direction than instructed, and 100 points when the specified limit was reached. These points were accumulated after each regulation block, and the total was presented to the participant in the end of each. After each run, participants were asked to verbally rate their degree of control over the disc size as well as the difficulty of the regulation in either direction on a scale from 0 to 10. In between runs, participants could take a short break (inside the scanner), if needed. At the end of the session and outside the scanner, participants were asked to write down the strategies they used, if they thought these strategies worked and whether they would use them again in the next session.

#### 2.4.3 Session 5: Transfer

After the standardized stimulation intensity adjustment an anatomical image was recorded and the eye tracker was calibrated. The subsequent three functional runs were similar to the feedback runs in previous sessions, but differed in two ways: (1) no feedback was given to the participant about their regulation performance, and (2) during 50% of all rest and regulation blocks, there was the chance (11.8% across all three runs) of a mild electrical stimulation of the fingers at any time during the block. Participants were told to apply those strategies to increase and decrease the size of the disc that they thought worked best during the training, and that they were still collecting points. In the last run of the session a functional resting-state data-set was recorded while participants were asked to look at a fixation cross at the centre of the screen. At the end of the session and outside the scanner, participants were asked about any thoughts they would like to share about their participation, after which they were debriefed and informed about the details of the study.

### 2.5 Data analysis

#### 2.5.1 Self-evaluation

Perceived controllability of the feedback signal as well as perceived difficulty to control the signal in either direction specifically were assessed for each training session by averaging the scores across all runs of that session. Expected increase in perceived controllability from first to last training session, as well as decreased perceived difficulty in both regulation directions, were each tested for with a one-sided paired *t*-test. Analyses were performed using Pingouin (version 0.3.8; Vallat, 2018).

#### 2.5.2 Peripheral recordings

Bandpass filtering (0.3 – 3 Hz) was applied to raw pulse and respiratory recordings to remove low-frequency drifts, artefacts time-locked to the MR volumes were removed from the data via deconvolution, and automatic peak detection was applied to the pulse data, using a custom tool (https://github.com/can-lab/brainampconverter). Data was further visually inspected and corrected, using a custom tool (https://github.com/can-lab/hera). Periods rejected due to data quality were removed and data were interpolated. The processed pulse and respiratory data were then used for retrospective image-based correction of physiological noise artefacts in the MRI data, using a custom tool (https://github.com/can-lab/RETROICORplus). This method utilises 5th order Fourier modelling of cardiac and respiratory phase related noise. A total of 25 nuisance regressors were created, including 10 cardiac phase regressors and 10 respiratory phase regressors (RETROICOR; Glover et al., 2000), plus 3 heart rate frequency regressors (Shmueli et al., 2007; van Buuren et al,. 2009), and 2 respiratory volume per unit time regressors (Birn et al., 2006; van Buuren et al., 2009).

Skin conductance data recorded during the Transfer session were down-sampled to 100 Hz and high-pass filtered (cutoff = 5 Hz). Continuous Decomposition Analysis on the first 10 seconds of each block was then performed in Ledalab (Benedek & Kaernbach, 2010), in order to extract tonic and phasic components of the skin conductance amplitude. To test whether the threat of a mild electric shock led to an expected overall increase in skin conductance, compared to not receiving a shock, the standardized (z-transformed) average phasic amplitude of the initial ten seconds of each block was calculated and subsequently entered into a one-sided paired t-test. In case of a significant overall effect, the expected increase was further assessed by post-hoc one-sided paired t-tests in each condition (rest, regulate to SN [participants 3, 5, 7, 9, 11: “larger”; participants 2, 4, 6, 8, 10: “smaller”], regulate to ECN [participants 3, 5, 7, 9, 11: “smaller”; participants 2, 4, 6, 8, 10: “larger”]) individually. Potential differences in the strength of the threat effect between conditions was furthermore tested by the interaction effect in a 3 × 2 repeated measures ANOVA with the factors condition (rest, regulate to SN, regulate to ECN) and threat (threat, safe).

Average baseline pupil dilation in response to each stimulus was calculated from data acquired in the Localiser session. The average pupil size of the first 10 seconds of each rest and regulation block was then calculated for the Transfer session and z-transformed, from which the z-transformed average baseline was subtracted, to avoid pupil dilation differences driven by changes in luminance between stimuli. Periods reported as blinks by the eye-tracker (pupil data missing for three or more samples in a sequence) were not considered. To test whether the threat of a mild electric shock led to an overall increase in pupil size, compared to not receiving a shock, baseline-corrected data was subsequently entered into a one-sided paired t-test. In case of a significant overall effect, the expected increase was further assessed by post-hoc one-sided paired t-tests in each condition (rest, regulate to SN [participants 3, 5, 7, 9, 11: “larger”; participants 2, 4, 6, 8, 10: “smaller”], regulate to ECN [participants 3, 5, 7, 9, 11: “smaller”; participants 2, 4, 6, 8, 10: “larger”]) individually. Potential differences in the strength of the threat effect between conditions was furthermore tested by the interaction effect in a 3 × 2 repeated measures ANOVA with the factors condition (rest, regulate to SN, regulate to ECN) and threat (threat, safe). Skin conductance and pupil size analyses were performed using Pingouin (version 0.3.8; Vallat, 2018).

#### 2.5.3 SN-ECN balance self-regulation performance

The first two runs of the first training session were considered to be practice runs (for the participants to get used to the setting) and were not analysed further. For each remaining functional run, the difference signal between SN and ECN timecourses (as extracted online with Turbo-BrainVoyager) was high-pass filtered (cutoff = 0.01 Hz/100 s) and normalized (z-transformation). For each participant, the concatenated preprocessed difference signals of all runs of all sessions entered a generalized linear model (GLM), corrected for serial correlations by means of a first order autoregressive model (AR1), with 29 regressors modelling the expected hemodynamic responses (double gamma function) during the different blocks of the experimental design in each session (i.e. the regulation conditions “larger” and “smaller”, the “delay” period, and “feedback” for each regulation condition in each of the training sessions, as well as the regulation conditions “larger”, “smaller”, “larger|threat”, “smaller|threat”, “rest|threat”, and regressors for reinforced threat blocks describing the periods before and after a shock, as well as the shock itself, for the transfer session). An additional set of 49 regressors per run was added as covariates: six motion parameters from Turbo-BrainVoyager (three translational and three rotational), their first temporal derivative as well as the quadratic terms of both the base motion parameters and their temporal derivatives, and 25 physiological noise components. Self-regulation performance in each session was assessed by estimating the difference contrast “regulate to SN > regulate to ECN” (participants 3, 5, 7, 9, 11: “larger > smaller”; participants 2, 4, 6, 8, 10: “smaller > larger”) corresponding to that session. In line with our hypotheses, the following planned contrasts were tested for significance: (1) the difference contrast across all sessions, to test for overall control over the feedback signal, (2) the difference contrast in the transfer session, to test specifically for preserving that skill after training, in the absence of feedback, (3) a contrast corresponding to a positive linear trend in the difference contrast over sessions, to test for improvement over time, and (4) a contrast testing whether the effect of the difference contrast in the transfer session was smaller in the “threat” condition, compared to the “safe” condition. To assess random effects across participants, contrast estimates of all participants where tested for significance with a one-sample *t*-test. The significance level for all tests was set to *α* = 0.05. Analyses were performed using NiPy (version 0.4.2; Millman, 2007) and Pingouin (version 0.3.8; Vallat, 2018).

#### 2.5.4 Whole-brain voxel-wise analysis

To further explore how the self-regulation of SN-ECN balance affected global brain activations in the here tested group of participants, additional whole-brain voxel-wise offline fMRI analysis has been performed. All MR image were preprocessed with FMRIPREP version 1.5.8 (Esteban et al., 2018; Esteban et al. 2020; RRID:SCR_016216), a Nipype (Gorgolewski et al., 2011; Gorgolewski et al., 2017; RRID:SCR_002502) based tool. Each T1w (T1-weighted) volume was corrected for INU (intensity non-uniformity) using `N4BiasFieldCorrection` v2.1.0 (Tustison et al., 2010) and skull-stripped using `antsBrainExtraction.sh` v2.1.0 (using the OASIS template). Spatial normalization to the ICBM 152 Nonlinear Asymmetrical template version 2009c (Fonov et al., 2009; RRID:SCR_008796) was performed through nonlinear registration with the `antsRegistration` tool of ANTs v2.1.0 (Avants et al., 2008; RRID:SCR_004757), using brain-extracted versions of both T1w volume and template. Brain tissue segmentation of cerebrospinal fluid (CSF), white-matter (WM) and gray-matter (GM) was performed on the brain-extracted T1w using fast (Zhang et al, 2001; FSL v5.0.9; RRID:SCR_002823).

Functional data was motion corrected using `mcflirt` (FSL v5.0.9; Jenkinson et al., 2002). This was followed by co-registration to the corresponding T1w using boundary-based registration (Greve et al., 2009) with six degrees of freedom, using `flirt` (FSL). Motion correcting transformations, BOLD-to-T1w transformation and T1w-to-template (MNI) warp were concatenated and applied in a single step using `antsApplyTransforms` (ANTs v2.1.0) using Lanczos interpolation.

Many internal operations of FMRIPREP use Nilearn (Abraham et al., 2014; RRID:SCR_001362), principally within the BOLD-processing workflow. For more details of the pipeline see https://fmriprep.readthedocs.io/en/latest/workflows.html.

FMRIPREP-preprocessed functional runs were further spatially smoothed (5 mm FWHM) and temporally high-pass filtered (cutoff = 0.01 Hz/100 s), using a custom-made Nipype (version 1.4.2; Gorgolewski et al, 2011; Gorgolewski et al., 2017) pipeline (https://github.com/can-lab/finish-the-job).

The first two runs of the first training session were considered to be practice runs (for the participants to get used to the setting) and were not analysed further. Each of the remaining runs was median-scaled to a value of 10000, and all voxels exceeding a threshold of 1000 entered a pre-whitened generalized linear model (GLM; `FILM_GLS` from FSL version 6.0.1; Smith et al., 2004) with five regressors modelling the expected hemodynamic responses (double gamma function) during the different blocks of the experimental design (the regulation conditions “larger” and “smaller”, the “delay” period, as well as “feedback” for each regulation condition). An additional set of 49 regressors per run was added as covariates: 24 motion parameters from FMRIPREP (three translational and three rotational, their first temporal derivative, as well as the quadratic terms of both base motion parameters and their derivatives) and 25 physiological noise components from RETROICOR. First-level contrasts corresponding to “regulate to SN” (participants 3, 5, 7, 9, 11: “larger”; participants 2, 4, 6, 8, 10: “smaller”), “regulate to ECN” (participants 3, 5, 7, 9, 11: “smaller”; participants 2, 4, 6, 8, 10: “larger”), and “regulate to SN > regulate to ECN” (participants 3, 5, 7, 9, 11: “larger > smaller”; participants 2, 4, 6, 8, 10: “smaller > larger”), were estimated and averaged across all runs of one training session in a single-regressor second/intermediate-level fixed-effects GLM (`FLAMEO` from FSL; version 6.0.1; Smith et al., 2004). The transfer session was analysed in the same way, with additional regressors for the conditions “larger|threat”, “smaller|threat”, “rest|threat”, and regressors for reinforced threat blocks describing the periods before and after a shock, as well as the shock itself, for the transfer session, and additional contrasts for each regulation direction and the difference between them were estimated for safe and threat conditions independently. Session estimates then entered several third/group-level fixed-effects GLMs (`FLAMEO` from FSL version 6.0.1; Smith et al., 2004): four single-regressor models to test for main effects of each session, (2) a model with one regressors describing a positive linear relationship across sessions and 10 regressors describing subject effects (to test for improvements over time), as well as (3) a model with one regressor describing the difference between safe and threat conditions in the transfer session and 10 regressors describing subject effects. Results were family-wise error (FWE) corrected for multiple comparisons on the single voxel level, and thresholded at a *α* = 0.05. Analyses were performed using a custom-made Nipype (version 1.4.2; Gorgolewski et al, 2011; Gorgolewski et al., 2017) pipeline (https://github.com/can-lab/FawN).

#### 2.5.5 Data and code availability

Scripts used during real-time neurofeedback training, as well as scripts for reproducing the reported self-regulation performance, self-evaluation, peripheral recordings, and full-brain analyses are openly available in the Open Science Framework at https://osf.io/sh2ck. Pseudonymized data will be available on request from the Donders Repository at https://data.donders.ru.nl. Raw MR images are not publicly available due to privacy or ethical restrictions.

## 3 Results

### 3.1 SN-ECN balance self-regulation performance

Figure 4 shows self-regulation performance per session for each participant individually (top) as well as for the group of all participants (bottom). Most importantly, and in line with our hypothesis, participants gained significant differential control (i.e. between the two regulation directions) of the feedback signal, both, across all sessions, *t*(9) = 4.78, *p* < 0.001, *d* = 1.51, as well as during the transfer session, without feedback, in particular, *t*(9) = 3.38, *p* < 0.01, *d* = 1.07. This skill was even significantly demonstrable on an individual level for 8/10 and 7/10 participants, respectively (see Supplementary Material for detailed individual results). Participants furthermore showed a linear improvement in self-regulation performance over time, from the first training session to the transfer session, *t*(9) = 2.82, *p* < 0.05, *d* = 0.89. No significant difference between differential self-regulation performance in the “safe”, compared to the “threat” condition of the transfer session could be observed, *t*(9) = 1.05, *p* = 0.16, *d* = 0.33, and confidence intervals indicate significant differential control in both conditions. For a full overview of individuals’ self-regulation performance see Supplementary Materials.

**Figure 3.**
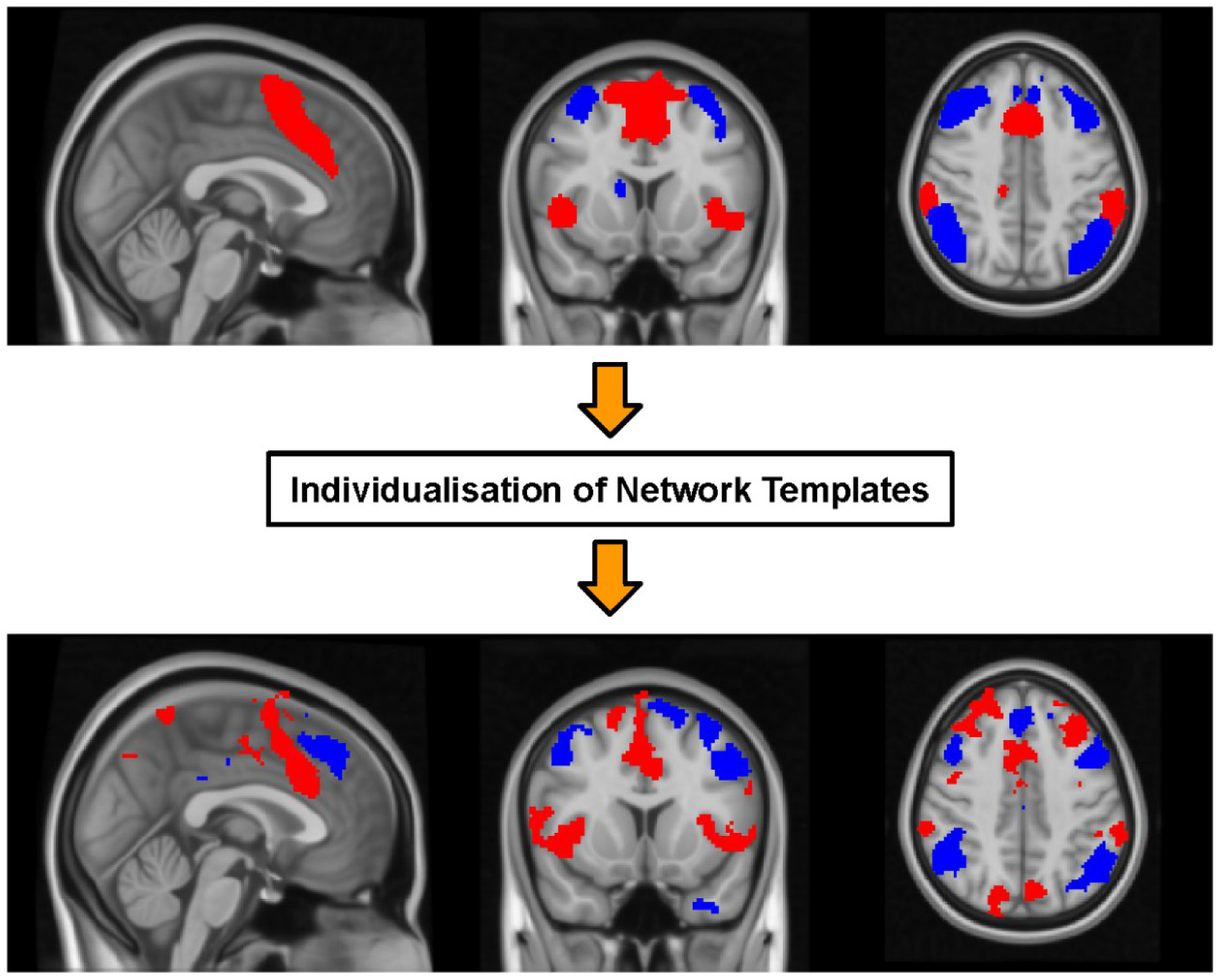
Example output of procedure to individualise SN (red) and ECN (blue) network templates (top) to subject-specific network masks (bottom; data from participant 11, back-projected from native space to MNI space).

**Figure 4.**
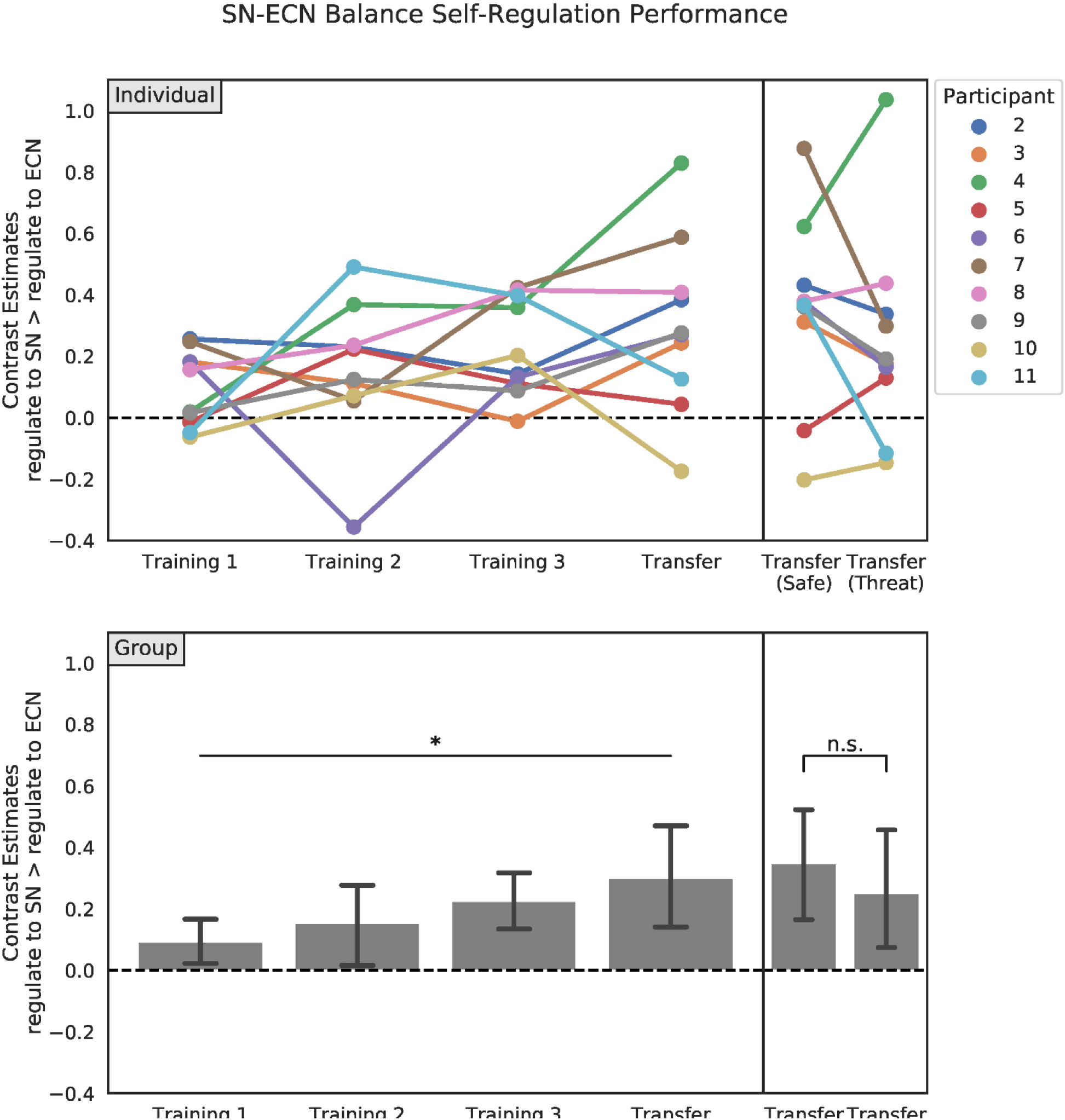
SN-ECN balance self-regulation performance per session for each participant individually (top) and for all participants as a group (bottom). On average, participants gained significant control over the feedback signal, irrespective of being threatened or not, and demonstrated consistent improvement during acquisation of this skill. Error bars represent 95% bootstrapped confidence intervalls, and indicate significance when they do not encompass zero. **: p < 0.01, n.s.: not significant.

### 3.2 Self-evaluation

There was a significant increase in the ratings of perceived control of the feedback signal from the first training session (3.91) to the last (4.71), *t*(9) = 2.06, *p* < 0.05, *d* = 0.42. A significant decrease in the rating of perceived regulation difficulty was observed when regulating to ECN (first training session: 7.66, last training session: 5.72), *t*(9) = −4.68, *p* < 0.001, *d* = 1.08, but not when regulating to SN (first training session: 5.77, last training session: 6.21), *t*(9) = 0.90, *p* = 0.80, *d* = 0.34. Participants’ strategies to regulate to SN included focusing on emotional memories/thoughts (n = 5), imagery of size changes (n = 3), as well as refocusing covert attention away from the task (n = 2). Strategies to regulate to ECN included focusing on positive memories/relaxing thoughts (n = 5), imagery of size changes (n = 3), as well as mental calculation (n = 2). For a full overview of individuals’ ratings and strategies per session see Supplementary Materials.

### 3.3 Physiological threat response

Figure 5 shows pupil responses in the transfer runs. Overall, normalized pupil size showed the expected sympathetic response to the stressor, *t*(9) = 3.16, *p* < 0.01, *d* = 1.89, with larger pupil size during threat blocks (0.42 SD) than during safe blocks (−0.34 SD), validating the experimental manipulation. Post-hoc tests indicated this effect to be significant during both rest, *t*(9) = 3.21, *p* < 0.01, *d* = 1.77, and when regulating towards ECN, *t*(9) = 2.13, *p* < 0.05, *d* = 1.04, but not when regulating towards SN, *t*(9) = 0.82, *p* = 0.23, *d* = 0.46. The difference in the effect between conditions was not significant, *F*(2,18) = 3.45, *p* = 0.054, *η_p_^2^* = 0.28.

**Figure 5.**
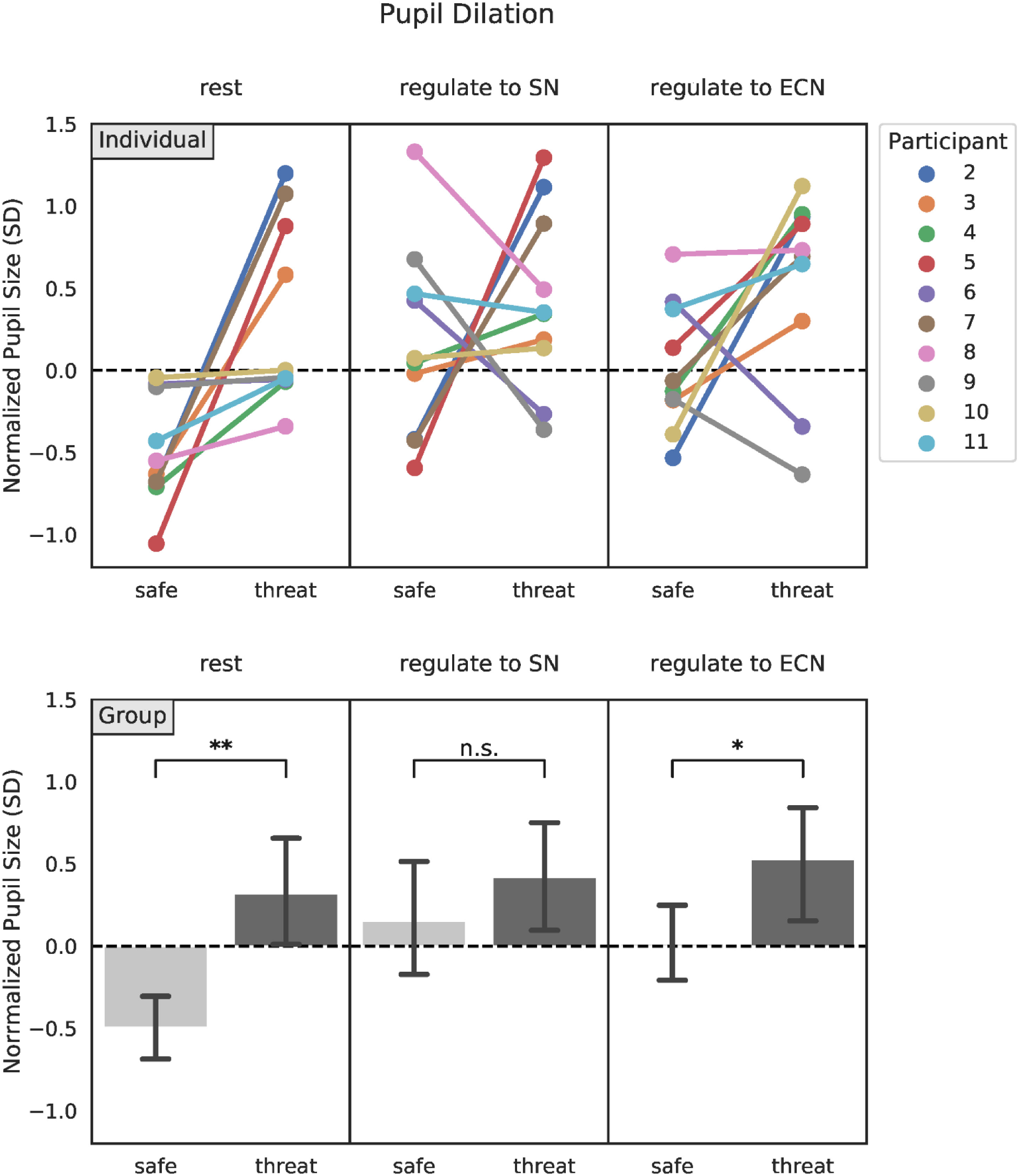
Pupil dilation in the Transfer session for each participant individually (top) and for all participants as a group (bottom). The threat of a mild electric stimlation significantly increased overall pupil size, but to different degrees in each task. Error bars represent 95% confidence intervalls. **: *p* < 0.01, *: *p* < 0.05, n.s.: not significant.

Overall, the same pattern was observed for standardized average phasic skin conductance amplitude in the transfer runs, which was 0.14 SD larger during threat blocks than during safe blocks, *t*(9) = 2.17, *p* < 0.05, *d* = 0.99. Post-hoc tests indicated that this effect reached significant only when regulating towards ECN, *t*(9) = 2.72, *p* < 0.05, *d* = 1.30, but not when regulating towards SN, *t*(9) = 1.39, *p* = 0.10, *d* = 0.70, or during rest, *t*(9) = 1.74, *p* = 0.06, *d* = 0.73. The difference in the effect between conditions was not significant, *F*(2,18) = 0.38, *p* = 0.69, *η_p_^2^* = 0.04.

### 3.4 Whole-brain voxel-wise analysis

Figure 6 shows an overview of the whole-brain voxel-wise effects of self-regulation in each of the four sessions, corrected for family-wise error. Regulating SN-ECN balance to either side positively activated a large set of regions associated with SN (including supplementary motor area, anterior cingulate cortex, supramarginal gyrus, insula, amygdala, hippocampus, dorsal striatum, thalamus), and negatively activated a large set of regions associated with ECN (including middle frontal gyrus, angular gyrus) and DMN (including posterior cingulate gyrus, precuneus, paracingulate gyrus). Descriptively, participants increasingly learned to differentially activate SN, but also part of DMN, between the two regulation conditions. Results of the contrast specifically testing for an improvement over time confirm this pattern, and further indicated that while participants did improve in increasingly activating SN when asked to regulate to SN, they mainly learned to actively deactivate SN when asked to shift SN-ECN balance in the other direction. Tables S1, S2 and S3 (Supplementary Materials) provide the full list of regions whose activation changed positively or negatively over time, for differential regulation, regulation to SN and regulation to ECN, respectively.

**Figure 6.**
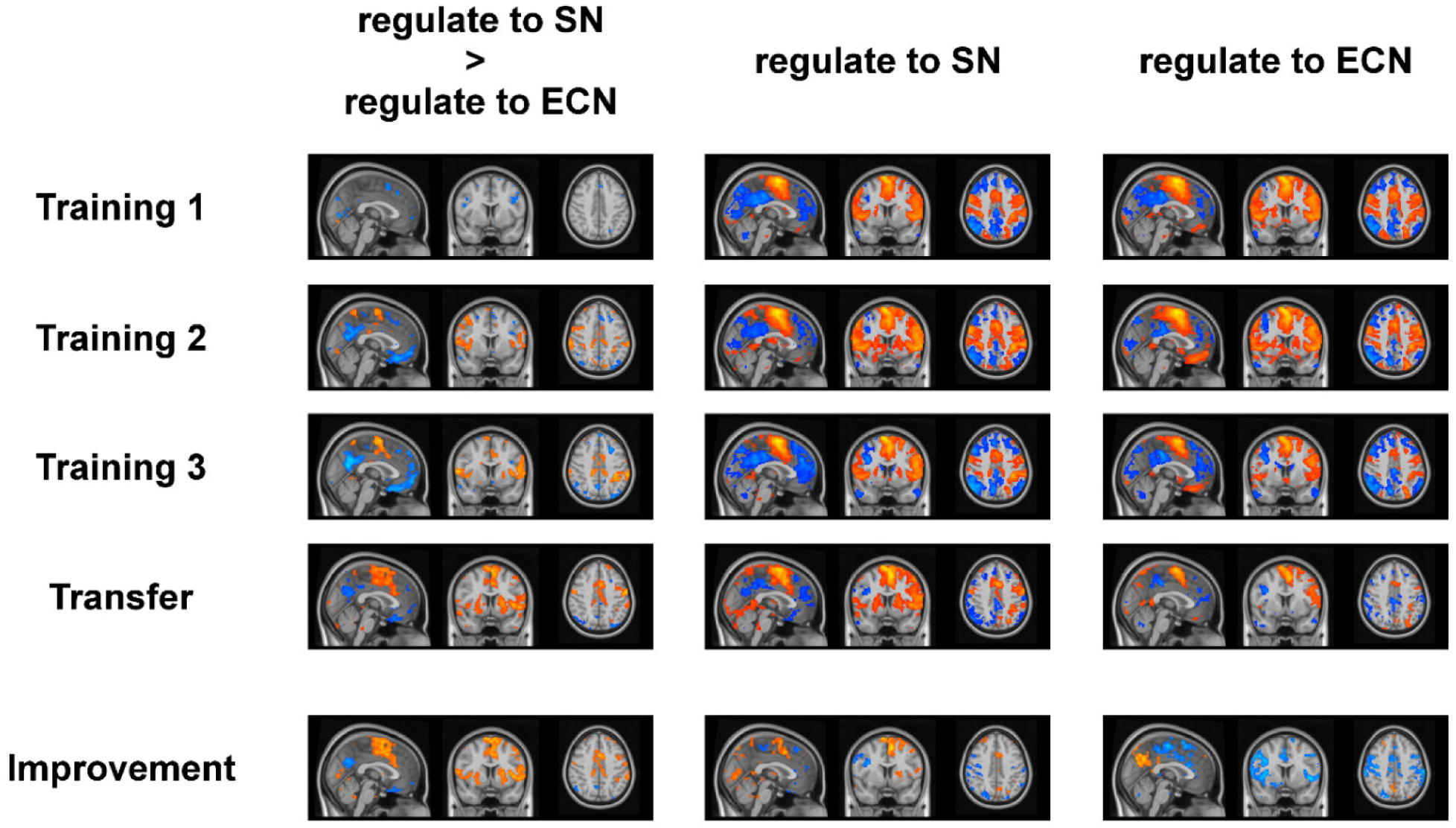
Full-brain activations of the tested group of participants during self-regulation in each of the four sessions. Participants learned to regulate the difference signal mainly via SN and ECN (but also DMN) manipulation, and especially by actively deactivating SN when asked to regulate to ECN.

Figure 7a shows the whole-brain voxel-wise effects of threat in the transfer session, corrected for family-wise error. Overall, being threatened with a mild electric stimulation led to an increase in activation of SN regions and visual brain areas, as well as a decrease in activation of ECN and DMN regions (see table S4 in Supplementary Materials for the full list of regions). This effect was more pronounced in the two regulation conditions, compared to the rest condition. Figure 7b shows the effects of threat on self-regulation in particular. Self-regulating to either direction, when being threatened with a mild electric shock, led to local increases in activation of visual brain areas, insular cortex, superior frontal gyrus and posterior cingulate cortex, compared to not being threatened (see tables S5 and S6 in Supplementary Materials for the full list of regions). However, in the differential contrast between the two regulation directions, only a single voxel in the middle frontal gyrus being was significantly more activated when being threatened, compared to when not being threatened.

**Figure 7.**
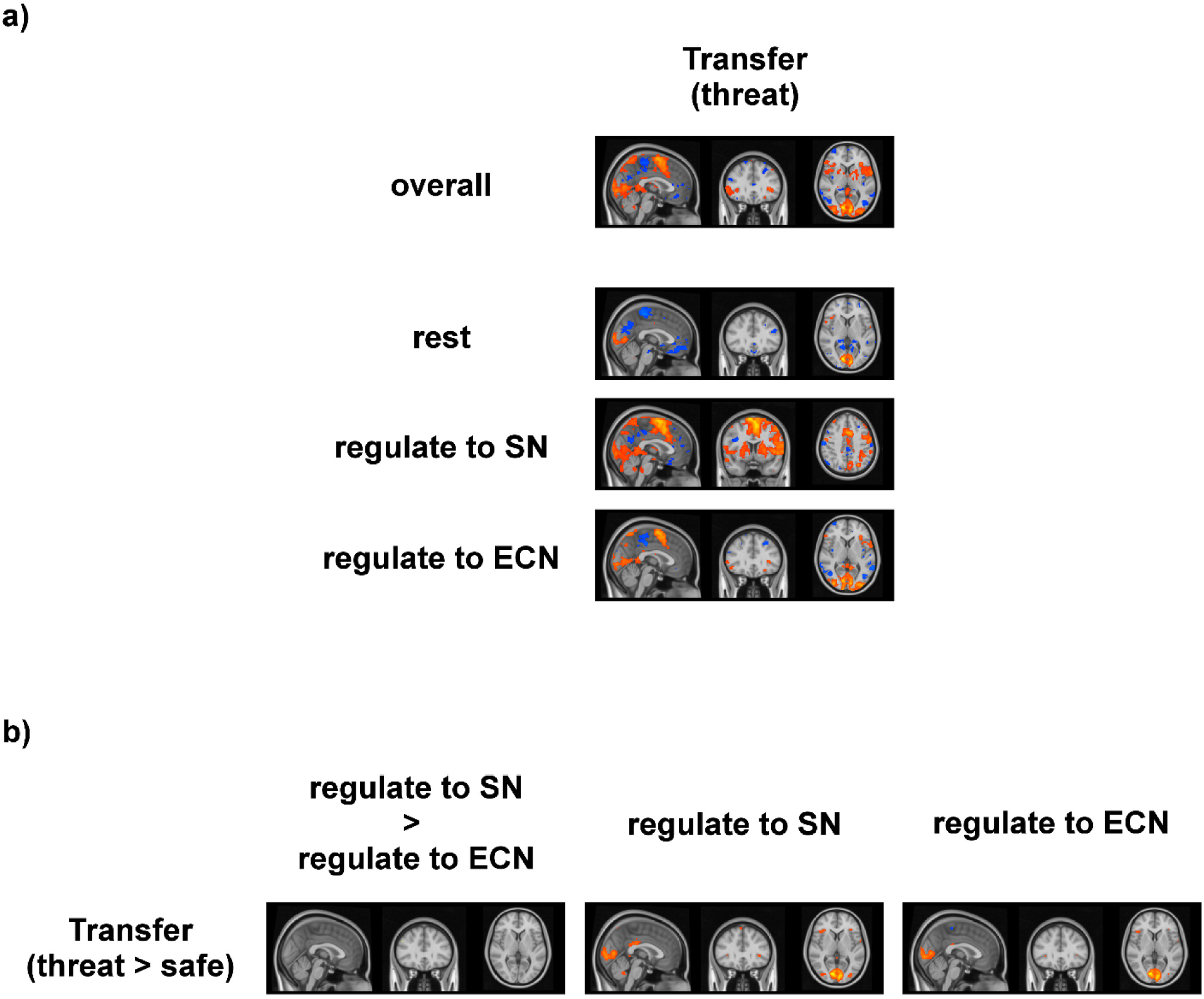
Full-brain activations during self-regulation in the transfer session. a) Being threatened with a mild electric shock led to an increase in SN activation, as well as decreases in ECN and DMN activation. (b) The threat affected self-regulation in both directions equally.

## 4 Discussion

The current study is a first investigation of the general feasibility to use rtfMRI-NF to train the self-regulation of stress-related large-scale network balance. Our results show that, given intermitted feedback about their performance over three training sessions, participants were well capable to learn to differentially control the balance between SN and ECN activation, both as a group, as well as individually.

Crucially, participants were able to successfully transfer this newly learned skill to a situation where they did not receive any feedback anymore. Unlike other forms of neuromodulation which externally change neural parameters at the time of application or aim for long-term effects via plasticity changes (Johnson et al., 2013), neurofeedback specifically allows individuals to learn mental strategies that they can voluntarily apply themselves at a later point in time after the training. While participants in the current study used very different mental strategies, ranging from emotion induction to exerting cognitive control, they were all able to explicitly describe their chosen strategies afterwards and seemed to be aware of their self-regulation success. In addition to transferring the learned self-regulation strategies beyond the training, participants were also capable to apply them under acute stress (in form of a threat of mild electric stimulation). The threat of mild electric stimulation led to a significant overall increase in both pupil size and skin conductance, compared to periods without this threat, validating that, in line with previous research (Phelps et al., 2001), the threat was perceived as an acute stressor. This effect, while descriptively present in all conditions, did not reach statistical significance in each of them individually (probably due to the small size of the current sample), but there was also no evidence that the effect significantly differed between conditions.

Whole-brain voxel-wise fMRI analysis confirmed that participants recruited large-scale networks of brain areas in SN and ECN, but also DMN, during their self-regulation attempts. Notably, participants’ attempts to regulate SN-ECN balance in either direction seem to have led to a general shift of network balance toward SN, with strong SN activation and ECN (and DMN) deactivation, compared to rest. We also consistently observed this pattern in the online feedback signal during the training sessions. There are multiple possible explanations for this observation. It has previously been shown that increases in cognitive effort as well as reward anticipation recruit overlapping networks of brain areas within SN (Vassena et al., 2014). Reward processing networks are known to be part of the neural substrates of neurofeedback-based self-regulation, together with other SN and ECN areas, involved in the conscious perception of feedback and reward, and executive aspects of the regulation tasks, respectively (Sitaram et al., 2017). Furthermore, an increase in cognitive effort during regulation attempts might have led to heightened sympathetic arousal (Westbrook & Braver, 2015), resulting in network balance shifts similar to acute stressors (Young et al., 2017). Alternatively, the pressure to perform self-regulation on cue within a limited time period might have simply been perceived as an acute stressor. The results of the threat manipulation in both whole-brain fMRI as well as pupil data seem to be in line with this interpretation. The pupil data in the transfer session showed visibly increased dilation during regulation blocks, compared to safe blocks (see Figure 5). The difference was more pronounced in the safe condition than in the threat condition, possibly due to pupil size ceiling effects when regulating under threat. Likewise, whole-brain voxel-wise fMRI data showed that while threat in general led to increases in SN activation and decreases in ECN and MN activation, these neural effects were largely driven by the regulation conditions (which appeared to be equally affected by the threat).

Importantly, however, despite a general shift towards SN, whole-brain voxel-wise fMRI analysis showed that participants were able to learn over time to suppress SN activation when regulating towards ECN, and continued to do so in the transfer session. This is also reflected in the self-evaluation results which indicated that participants, while generally aware of their improvement in self-regulation over time, only perceived a decrease in the difficulty to regulate to ECN over time. From a clinical perspective, the ability to voluntarily regulate network balance away from SN is particularly relevant, as it might allow patients with stress-related disorders to actively counteract the automatic stress-induced shifts towards SN (Hermans, et al., 2014). Future research applying this novel network-based neurofeedback approach to corresponding patient populations will be needed to test this hypothesis. It should be noted that, due to the explorative nature of the whole-brain voxel-wise fMRI analysis and the focus on understanding how the group of tested participants in the current study achieved self-regulation of the network balance, fixed effects analysis was performed. In contrast to the other analyses reported, the conclusions drawn from this analysis are hence not generalizable beyond the current sample.

The neurofeedback approach in the current study differs considerably from the majority of previous rtfMRI-NF paradigms, in that it does not target only the activation of a single isolated brain area (Thibault et al, 2017; Sulzer et al., 2013), nor the functional connectivity between two individual areas (Thibault et al, 2017; Watanabe et al, 2017). We specifically targeted the difference in activation between two functionally different actors (c.f. Scharnowski et al., 2015). Unlike Scharnowski and colleagues (2015), however, we focus on the balance between two large-scale brain networks that each consist of multiple brain areas and are functionally related. Our approach is in line with two other recent neurofeedback studies that target large-scale brain network balance (Kim et al. 2019; Pamplona et al., 2020). While Kim and colleagues (2019) targeted changes in functional connectivity between SN and DMN, Pamplona and colleagues (2020) took an approach similar to ours and targeted the difference in the activation between the sustained attention network and DMN. Notably, however, the current study differs to both of these studies by focusing on the balance between SN and ECN, specifically, providing another important proof-of-concept demonstration for a promising new class of network-based neurofeedback paradigms. Besides its clinical potential to target maladaptations in large-scale network configurations, network-based neurofeedback might also be less affected by physiological confounds (in particular respiratory patterns) that are otherwise difficult to remove from a single real-time neurofeedback signal (Weiss et al., 2020). The large-scale nature of the involved networks over large parts of the brain, combined with the characteristically widespread pattern in which physiological noise affects brain activity (Birn et al., 2006), suggest that this confounding factor would affect each network to a very similar degree, and that a subtraction of two of the signal from two of those networks would largely cancel out the noise. This, however, remains a theoretical argument at this point, which needs to be further detailed and specifically tested in future research. In the current study, we consequently applied physiological noise correction to the offline fMRI analyses to further mitigate this issue (Glover et al., 2000; Birn et al., 2006; Shmueli et al., 2007; van Buuren et al., 2009).

In conclusion, the current study constitutes an important first successful demonstration of neurofeedback training based on stress-related large-scale network balance – a novel approach that has the potential to train control over the central response to stressors in real-life situations outside of the MRI scanner, opening up new potential clinical approaches to changing maladaptive stress responses – the underlying mechanism of a large variety of mental disorders (de Kloet et al, 2005) – and to promote resilience (Kalisch et al, 2017; Kalisch et al., 2015).

## 5 Acknowledgments

This work was supported by a grant from the European Research Council (ERC-2015-CoG 682591). We would like to thank Rayyan Tutunji and Milette Dufour for their valuable input during the planning stage of this study.

## Footnotes

1 Notably, one individual (participant 6) scored unexpectedly high on the STAI-trait questionnaire (69). Excluding this participant from analysis, however, does not change the interpretation of the results or conclusions drawn in this study.

