## Supplementary Materials for "Self-regulation of stress-related large-scale brain network balance using real-time fMRI Neurofeedback"

The following pages include additional materials that were too comprehensive to be included in the published article.

For further supplements, please also see: https://osf.io/sh2ck.

### Individual self-regulation strategies and ratings

***Table 1.*** *Overview of self-regulation self-evaluation and strategies of each participant. 1 = control over signal, 2 = SN strategy & difficulty score, 3 = ECN strategy & difficulty score.*

| | | Training-1 | Score | Training-2 | Score | Training-3 | Score |
|---|---|---|---|---|---|---|---|
| | **1** | | 2,625 | | 5 | | 4,5 |
| sub 2 | **2** | thinking of small objects | 5,286 | thinking of small objects | 2,875 | thinking of small objects and deflation | 7 |
| | **3** | thinking of large objects | 8,714 | thinking of large objects | 8,75 | thinking of large objects and inflation | 6 |
| | **1** | | 0,875 | | 0,875 | | 1 |
| sub 3 | **2** | covert attention to the arrows | 6,857 | covert attention to the arrows | 9,625 | changing thoughts | 7,143 |
| | **3** | relaxing | 9 | formulating sentences in french | 0,25 | boring thoughts | 8,857 |
| | **1** | | 5,25 | | 7 | | 7 |
| sub 4 | **2** | sad feelings | 3,714 | sad feelings | 4,571 | sad feelings | 7,125 |
| | **3** | happy or angry feelings | 5,571 | happy memories | 5,571 | happy memories | 2,375 |
| | **1** | | 5,625 | | 5,75 | | 4,375 |
| sub 5 | **2** | get into a panic state | 5,857 | changing focus on surrounding sounds | 5,75 | memory task | 7,125 |
| | **3** | imagine a growing circle | 6,143 | counting, speaking different languages | 3,5 | counting | 5,5 |
| | **1** | | 4,125 | | 4,429 | | 5,5 |
| sub 6 | **2** | mean and aggressive thoughts | 5,571 | negative thoughts | 7,286 | negative and scary thoughts | 6 |
| | **3** | - | 7,714 | positive thoughts | 6,857 | complimented self and happy feelings | 5,75 |
| | **1** | | 4,75 | | 4,5 | | 6 |
| sub 7 | **2** | stressful memory | 5,429 | stressful memory | 3,75 | car-crash film memory | 4,571 |
| | **3** | calming memory | 8 | mind-wondering | 8 | calming memory | 4,429 |
| | **1** | | 6,5 | | 6,143 | | 6,875 |
| sub 8 | **2** | breaking items and being stressed | 6 | first- person actions (chores) | 5,286 | first- person actions (chores) | 4 |
| | **3** | positive thoughts | 7,571 | thinking of other people | 6 | thinking of other people | 4,875 |
| | **1** | | 2,375 | | 1,75 | | 1,5 |
| sub 9 | **2** | imagine a growing circle | 8 | imagine a growing circle | 8,625 | imagine a growing circle | 8,5 |
| | **3** | imagine a decreasing circle | 9,571 | imagine a decreasing circle | 8,625 | imagine a decreasing circle | 9,75 |
| | **1** | | 2,25 | | 6,375 | | 4,875 |
| sub 10 | **2** | imagine deflating circle | 5,143 | imagine to interact with the circle | 3,25 | imagine to interact with the circle | 5,875 |
| | **3** | imagine inflating circle | 6,143 | imagine to interact with the circle | 4,5 | stretching the circle | 4,25 |
| | **1** | | 4,75 | | 4,875 | | 5,5 |
| sub 11 | **2** | imagine inflating circle | 5,857 | positive memories | 4,875 | positive memories | 4,75 |
| | 3 | calculating backwards | 8,143 | calculating | 6,25 | negative thoughts | 5,375 |

### Offline fMRI analysis result tables

Clusters to be reported in result tables were defined as sets of minimally five continuous thresholded voxels, as well as their as sub-peaks with a minimal distance of 10 voxels (24 mm). Importantly, this was not done for statistical inference (all results are thresholded on voxel-level), but only to reduce the number of peaks that are not meaningfully different in terms of their location. Result tables were created with AltasReader (version 0.1.2; Notter et al., 2019).

***Table S1.*** *Regions whose activation changed positively or negatively over time for the contrast regulate to SN > regulate to ECN. X-, Y- and Z-coordinates are in MNI space; volume is in mm.*

| # | X | Y | Z | Z-STAT | VOLUME | AAL | HARVARD-OXFORD |
|---|---|---|---|---|---|---|---|
| **1** | -10 | 14 | 35 | 12,46 | 25468 | Cingulate_Mid_L | 39.0% Left_Cingulate_Gyrus_anterior_division; 25.0% Left_Paracingulate_Gyrus |
| **1** | -3 | -10 | 71 | 11,91 | 25468 | Supp_Motor_Area_L | 47.0% Left_Juxtapositional_Lobule_Cortex_(formerly_Supplementary_Motor_Cortex); 11.0% Left_Precentral_Gyrus |
| **1** | -1 | -8 | 47 | 9,00 | 25468 | Cingulate_Mid_L | 50.0% Left_Juxtapositional_Lobule_Cortex_(formerly_Supplementary_Motor_Cortex); 32.0% Left_Cingulate_Gyrus_anterior_division |
| **1** | -15 | -34 | 42 | 7,70 | 25468 | Cingulate_Mid_L | 35.0% Left_Precentral_Gyrus; 13.0% Left_Cingulate_Gyrus_posterior_division; 10.0% Left_Postcentral_Gyrus; 9.0% Left_Precuneous_Cortex |
| **1** | -3 | 23 | 61 | 6,98 | 25468 | Supp_Motor_Area_L | 78.0% Left_Superior_Frontal_Gyrus |
| **2** | -44 | 9 | 1 | 11,10 | 13431 | Insula_L | 40.0% Left_Frontal_Operculum_Cortex; 33.0% Left_Central_Opercular_Cortex; 7.0% Left_Insular_Cortex |
| **2** | -58 | 14 | 25 | 9,37 | 13431 | Frontal_Inf_Oper_L | 33.0% Left_Inferior_Frontal_Gyrus_pars_opercularis; 14.0% Left_Precentral_Gyrus |
| **2** | -48 | 35 | 1 | 9,11 | 13431 | Frontal_Inf_Tri_L | 40.0% Left_Inferior_Frontal_Gyrus_pars_triangularis; 25.0% Left_Frontal_Pole |
| **3** | 35 | -51 | -28 | 12,26 | 8362 | Cerebelum_6_R | 0% no_label |
| **3** | 19 | -70 | -28 | 8,72 | 8362 | Cerebelum_6_R | 0% no_label |
| **4** | -58 | -41 | 32 | 14,36 | 6326 | SupraMarginal_L | 39.0% Left_Supramarginal_Gyrus_posterior_division; 34.0% Left_Supramarginal_Gyrus_anterior_division; 7.0% Left_Planum_Temporale |
| **5** | -22 | 4 | 8 | 9,70 | 6135 | Putamen_L | 95.0% Left_Putamen |
| **5** | -5 | -20 | -1 | 5,88 | 6135 | Thalamus_L | 80.0% Left_Thalamus |
| **6** | 54 | 33 | 1 | 9,42 | 6080 | Frontal_Inf_Tri_R | 51.0% Right_Inferior_Frontal_Gyrus_pars_triangularis; 31.0% Right_Frontal_Pole |
| **6** | 59 | 6 | 11 | 9,34 | 6080 | Rolandic_Oper_R | 55.0% Right_Precentral_Gyrus; 9.0% Right_Inferior_Frontal_Gyrus_pars_opercularis |
| **6** | 33 | 16 | 11 | 7,50 | 6080 | Insula_R | 41.0% Right_Frontal_Operculum_Cortex; 27.0% Right_Insular_Cortex |
| **7** | -39 | -1 | 59 | 11,21 | 5014 | Precentral_L | 51.0% Left_Middle_Frontal_Gyrus; 28.0% Left_Precentral_Gyrus |
| **8** | -29 | -60 | -23 | 9,49 | 4782 | Cerebelum_6_L | 0% no_label |
| **9** | -39 | 37 | 35 | 10,59 | 3839 | Frontal_Mid_2_L | 54.0% Left_Middle_Frontal_Gyrus; 35.0% Left_Frontal_Pole |
| **9** | -20 | 47 | 47 | 6,16 | 3839 | Frontal_Sup_2_L | 11.0% Left_Frontal_Pole |
| **10** | 64 | -29 | 23 | 10,32 | 3812 | Temporal_Sup_R | 31.0% Right_Parietal_Operculum_Cortex; 26.0% Right_Supramarginal_Gyrus_anterior_division; 12.0% Right_Planum_Temporale |
| **11** | -1 | -63 | 30 | -9,33 | 3416 | Precuneus_L | 88.0% Left_Precuneous_Cortex |
| **12** | -53 | -44 | 8 | 8,95 | 2582 | Temporal_Mid_L | 26.0% Left_Supramarginal_Gyrus_posterior_division; 15.0% Left_Middle_Temporal_Gyrus_temporooccipital_part; 12.0% Left_Superior_Temporal_Gyrus_posterior_division; 5.0% Left_Middle_Temporal_Gyrus_posterior_division |
| **13** | 23 | 2 | 8 | 7,06 | 2323 | Putamen_R | 94.0% Right_Putamen |
| **14** | 14 | -41 | 54 | 8,28 | 1981 | Paracentral_Lobule_R | 23.0% Right_Postcentral_Gyrus; 22.0% Right_Precuneous_Cortex |
| **14** | 11 | -22 | 40 | 7,73 | 1981 | Cingulate_Mid_R | 59.0% Right_Cingulate_Gyrus_posterior_division; 7.0% Right_Precentral_Gyrus |

| | | | | | | | |
|---|---|---|---|---|---|---|---|
| **15** | -44 | -65 | 35 | -7,62 | 1899 | Angular_L | 68.0% Left_Lateral_Occipital_Cortex_superior_division; 6.0% Left_Angular_Gyrus |
| **16** | 35 | 45 | 35 | 9,76 | 1708 | Frontal_Mid_2_R | 78.0% Right_Frontal_Pole |
| **17** | 45 | -1 | 54 | 10,49 | 1667 | Frontal_Mid_2_R | 46.0% Right_Precentral_Gyrus; 20.0% Right_Middle_Frontal_Gyrus |
| **18** | 52 | -8 | -18 | -7,47 | 1257 | Temporal_Mid_R | 27.0% Right_Middle_Temporal_Gyrus_posterior_division; 18.0% Right_Middle_Temporal_Gyrus_anterior_division; 11.0% Right_Superior_Temporal_Gyrus_anterior_division; 9.0% Right_Superior_Temporal_Gyrus_posterior_division |
| **19** | -65 | -6 | -18 | -8,50 | 1230 | Temporal_Mid_L | 45.0% Left_Middle_Temporal_Gyrus_anterior_division; 17.0% Left_Middle_Temporal_Gyrus_posterior_division |
| **20** | -53 | -60 | 6 | 11,18 | 1093 | Temporal_Mid_L | 36.0% Left_Middle_Temporal_Gyrus_temporooccipital_part; 20.0% Left_Lateral_Occipital_Cortex_inferior_division; 6.0% Left_Angular_Gyrus |
| **21** | 52 | -25 | 1 | 9,48 | 1079 | Temporal_Sup_R | 26.0% Right_Superior_Temporal_Gyrus_posterior_division |
| **22** | 42 | -70 | 49 | -8,09 | 1066 | Angular_R | 57.0% Right_Lateral_Occipital_Cortex_superior_division |
| **23** | -1 | 11 | -25 | -7,27 | 888 | no_label | 0% no_label |
| **24** | 7 | 45 | 30 | -7,46 | 765 | Cingulate_Mid_R | 46.0% Right_Paracingulate_Gyrus; 21.0% Right_Superior_Frontal_Gyrus |
| **25** | -24 | -51 | 68 | 6,81 | 656 | Parietal_Sup_L | 57.0% Left_Superior_Parietal_Lobule; 5.0% Left_Postcentral_Gyrus |
| **26** | -36 | 64 | 6 | 8,22 | 642 | no_label | 21.0% Left_Frontal_Pole |
| **27** | 23 | -99 | -4 | 7,55 | 560 | Occipital_Inf_R | 78.0% Right_Occipital_Pole |
| **28** | -41 | -3 | -11 | 7,52 | 533 | Temporal_Sup_L | 67.0% Left_Insular_Cortex |
| **29** | 42 | -56 | 20 | -7,73 | 465 | Temporal_Mid_R | 26.0% Right_Angular_Gyrus; 6.0% Right_Middle_Temporal_Gyrus_temporooccipital_part |
| **30** | -24 | -13 | -20 | -6,31 | 451 | Hippocampus_L | 95.0% Left_Hippocampus |
| **31** | 23 | 25 | 56 | -7,47 | 451 | Frontal_Sup_2_R | 65.0% Right_Superior_Frontal_Gyrus |
| **32** | 40 | 56 | 4 | -6,47 | 424 | Frontal_Mid_2_R | 93.0% Right_Frontal_Pole |
| **33** | -22 | -29 | 64 | 7,30 | 424 | Postcentral_L | 33.0% Left_Postcentral_Gyrus; 32.0% Left_Precentral_Gyrus |
| **34** | -29 | -80 | 47 | -6,79 | 410 | Parietal_Inf_L | 64.0% Left_Lateral_Occipital_Cortex_superior_division |
| **35** | -8 | -51 | 61 | 7,42 | 396 | Precuneus_L | 46.0% Left_Precuneous_Cortex; 18.0% Left_Postcentral_Gyrus; 5.0% Left_Superior_Parietal_Lobule |
| **36** | -24 | -60 | 1 | 6,44 | 383 | no_label | 15.0% Left_Lingual_Gyrus; 6.0% Left_Precuneous_Cortex |
| **37** | 19 | -29 | 61 | 7,44 | 383 | Precentral_R | 39.0% Right_Precentral_Gyrus; 26.0% Right_Postcentral_Gyrus |
| **38** | -65 | -22 | -1 | 7,74 | 369 | Temporal_Mid_L | 42.0% Left_Superior_Temporal_Gyrus_posterior_division; 10.0% Left_Middle_Temporal_Gyrus_posterior_division |
| **39** | -1 | -80 | 4 | 7,32 | 355 | Lingual_L | 33.0% Left_Lingual_Gyrus; 32.0% Left_Intracalcarine_Cortex; 7.0% Left_Supracalcarine_Cortex; 6.0% Right_Lingual_Gyrus; 5.0% Right_Intracalcarine_Cortex |
| **40** | 4 | -51 | -32 | 6,21 | 342 | Vermis_10 | 0% no_label |
| **41** | -3 | -56 | -1 | 6,15 | 342 | Vermis_4_5 | 6.0% Left_Lingual_Gyrus |
| **42** | 66 | -34 | -11 | -6,41 | 342 | Temporal_Mid_R | 67.0% Right_Middle_Temporal_Gyrus_posterior_division; 9.0% Right_Middle_Temporal_Gyrus_temporooccipital_part |
| **43** | 30 | -44 | 59 | 6,99 | 314 | Postcentral_R | 53.0% Right_Superior_Parietal_Lobule; 11.0% Right_Postcentral_Gyrus |
| **44** | -15 | -65 | 28 | -7,25 | 314 | Cuneus_L | 37.0% Left_Precuneous_Cortex; 7.0% Left_Supracalcarine_Cortex; 6.0% Left_Cuneal_Cortex |
| **45** | -39 | -44 | 42 | 6,81 | 314 | Parietal_Inf_L | 29.0% Left_Superior_Parietal_Lobule; 28.0% Left_Supramarginal_Gyrus_posterior_division; 6.0% Left_Postcentral_Gyrus; 5.0% Left_Supramarginal_Gyrus_anterior_division |
| **46** | 4 | 59 | -4 | -6,85 | 314 | Frontal_Med_Orb_R | 86.0% Right_Frontal_Pole; 7.0% Right_Frontal_Medial_Cortex |
| **47** | -12 | -82 | 6 | 6,52 | 314 | Calcarine_L | 64.0% Left_Intracalcarine_Cortex |
| **48** | 30 | -10 | 4 | 6,71 | 273 | Putamen_R | 99.0% Right_Putamen |
| **49** | 47 | 37 | -16 | -6,18 | 273 | OFClat_R | 66.0% Right_Frontal_Pole; 18.0% Right_Frontal_Orbital_Cortex |
| **50** | 7 | -87 | 1 | 7,40 | 246 | Calcarine_R | 36.0% Right_Intracalcarine_Cortex; 21.0% Right_Lingual_Gyrus; 16.0% Right_Occipital_Pole |
| **51** | 9 | -65 | 11 | 5,96 | 232 | Calcarine_R | 62.0% Right_Intracalcarine_Cortex; 7.0% Right_Supracalcarine_Cortex |
| **52** | -5 | -75 | -18 | 6,74 | 219 | Cerebelum_6_L | 0% no_label |

| | | | | | | | |
|---|---|---|---|---|---|---|---|
| **53** | -44 | -63 | -1 | 6,39 | 205 | Temporal_Mid_L | 33.0% Left_Lateral_Occipital_Cortex_inferior_division; 20.0% Left_Middle_Temporal_Gyrus_temporooccipital_part; 9.0% Left_Inferior_Temporal_Gyrus_temporooccipital_part |
| **54** | -20 | 61 | 20 | -6,19 | 205 | Frontal_Sup_2_L | 80.0% Left_Frontal_Pole |
| **55** | -10 | 28 | -16 | -6,97 | 205 | Rectus_L | 8.0% Left_Subcallosal_Cortex |
| **56** | -27 | 35 | -16 | 7,11 | 178 | OFCant_L | 55.0% Left_Frontal_Orbital_Cortex; 39.0% Left_Frontal_Pole |
| **57** | -1 | -72 | 13 | 6,36 | 164 | Calcarine_L | 41.0% Left_Intracalcarine_Cortex; 26.0% Left_Supracalcarine_Cortex; 9.0% Left_Lingual_Gyrus |
| **58** | -48 | 28 | 37 | 6,50 | 164 | Frontal_Mid_2_L | 51.0% Left_Middle_Frontal_Gyrus |
| **59** | 9 | 56 | 6 | -7,04 | 150 | Frontal_Sup_Medial_R | 33.0% Right_Frontal_Pole; 26.0% Right_Paracingulate_Gyrus |
| **60** | -10 | -25 | 78 | 5,86 | 150 | Paracentral_Lobule_L | 51.0% Left_Precentral_Gyrus |
| **61** | 16 | -63 | 25 | -6,37 | 150 | Precuneus_R | 61.0% Right_Precuneous_Cortex; 13.0% Right_Supracalcarine_Cortex; 7.0% Right_Cuneal_Cortex |
| **62** | 26 | 61 | 23 | 6,33 | 150 | Frontal_Sup_2_R | 76.0% Right_Frontal_Pole |
| **63** | -29 | -99 | -4 | 6,21 | 137 | Occipital_Mid_L | 67.0% Left_Occipital_Pole |
| **64** | 19 | 40 | 54 | -6,59 | 123 | Frontal_Sup_2_R | 30.0% Right_Frontal_Pole; 13.0% Right_Superior_Frontal_Gyrus |
| **65** | 28 | -13 | 56 | 6,05 | 123 | Frontal_Sup_2_R | 48.0% Right_Precentral_Gyrus; 7.0% Right_Superior_Frontal_Gyrus |
| **66** | 16 | -8 | -18 | -6,17 | 109 | ParaHippocampal_R | 54.0% Right_Amygdala; 40.0% Right_Hippocampus |
| **67** | -12 | -70 | 6 | 5,90 | 96 | Calcarine_L | 50.0% Left_Intracalcarine_Cortex; 6.0% Left_Lingual_Gyrus |
| **68** | -27 | 21 | 37 | -6,13 | 96 | Frontal_Mid_2_L | 37.0% Left_Middle_Frontal_Gyrus |
| **69** | -10 | -39 | 37 | -6,23 | 96 | Cingulate_Mid_L | 31.0% Left_Cingulate_Gyrus_posterior_division |
| **70** | 26 | -10 | -23 | -6,06 | 96 | Hippocampus_R | 96.0% Right_Hippocampus |
| **71** | 14 | -6 | 16 | 5,61 | 82 | Caudate_R | 32.0% Right_Thalamus; 7.0% Right_Caudate |
| **72** | -51 | -25 | 40 | 5,65 | 82 | Parietal_Inf_L | 50.0% Left_Postcentral_Gyrus; 14.0% Left_Supramarginal_Gyrus_anterior_division |
| **73** | 2 | -13 | -35 | 6,63 | 82 | no_label | 32.0% Brain-Stem |
| **74** | 52 | -10 | -28 | -6,75 | 68 | Temporal_Inf_R | 20.0% Right_Middle_Temporal_Gyrus_posterior_division; 12.0% Right_Inferior_Temporal_Gyrus_posterior_division; 12.0% Right_Inferior_Temporal_Gyrus_anterior_division; 6.0% Right_Middle_Temporal_Gyrus_anterior_division |
| **75** | -29 | -44 | 71 | 5,67 | 68 | Parietal_Sup_L | 40.0% Left_Superior_Parietal_Lobule; 14.0% Left_Postcentral_Gyrus |
| **76** | -15 | -58 | 66 | 5,57 | 68 | Precuneus_L | 39.0% Left_Superior_Parietal_Lobule; 37.0% Left_Lateral_Occipital_Cortex_superior_division |
| **77** | 19 | -87 | -4 | -5,94 | 68 | Lingual_R | 19.0% Right_Lingual_Gyrus; 17.0% Right_Occipital_Pole; 13.0% Right_Occipital_Fusiform_Gyrus |
| **78** | -55 | -51 | 13 | -5,91 | 68 | Temporal_Mid_L | 33.0% Left_Angular_Gyrus; 25.0% Left_Supramarginal_Gyrus_posterior_division; 12.0% Left_Middle_Temporal_Gyrus_temporooccipital_part |
| **79** | -41 | -87 | 23 | -6,79 | 68 | Occipital_Mid_L | 47.0% Left_Lateral_Occipital_Cortex_superior_division; 12.0% Left_Occipital_Pole |

*__Table S2.__ Regions whose activation changed positively or negatively over time when regulating to SN. X-, Y- and Z-coordinates are in MNI space; volume is in mm.*

| # | X | Y | Z | Z-STAT | VOLUME | AAL | HARVARD-OXFORD |
|---|---|---|---|---|---|---|---|
| **1** | -3 | 4 | 61 | 15,90 | 20017 | Supp_Motor_Area_L | 81.0% Left_Juxtapositional_Lobule_Cortex_(formerly_Supplementary_Motor_Cortex); 10.0% Left_Superior_Frontal_Gyrus |
| **1** | -32 | -1 | 61 | 13,16 | 20017 | Precentral_L | 45.0% Left_Middle_Frontal_Gyrus; 10.0% Left_Superior_Frontal_Gyrus; 7.0% Left_Precentral_Gyrus |
| **1** | -10 | 14 | 35 | 11,39 | 20017 | Cingulate_Mid_L | 39.0% Left_Cingulate_Gyrus_anterior_division; 25.0% Left_Paracingulate_Gyrus |
| **1** | 23 | -3 | 66 | 7,35 | 20017 | Frontal_Sup_2_R | 47.0% Right_Superior_Frontal_Gyrus; 16.0% Right_Precentral_Gyrus |
| **2** | 61 | -1 | 35 | -13,35 | 13540 | Postcentral_R | 66.0% Right_Precentral_Gyrus; 16.0% Right_Postcentral_Gyrus |
| **2** | 59 | -29 | 16 | -9,69 | 13540 | Temporal_Sup_R | 50.0% Right_Planum_Temporale; 20.0% Right_Parietal_Operculum_Cortex |
| **2** | 38 | -10 | 18 | -9,49 | 13540 | Insula_R | 71.0% Right_Central_Opercular_Cortex; 8.0% Right_Insular_Cortex |
| **3** | 42 | -39 | 59 | -11,72 | 12707 | Postcentral_R | 41.0% Right_Superior_Parietal_Lobule; 17.0% Right_Postcentral_Gyrus; 8.0% Right_Supramarginal_Gyrus_posterior_division |
| **3** | 64 | -32 | 42 | -8,64 | 12707 | SupraMarginal_R | 41.0% Right_Supramarginal_Gyrus_anterior_division; 16.0% Right_Supramarginal_Gyrus_posterior_division; 5.0% Right_Parietal_Operculum_Cortex |
| **3** | 21 | -32 | 78 | -7,94 | 12707 | Postcentral_R | 31.0% Right_Postcentral_Gyrus |
| **3** | 23 | -56 | 71 | -7,40 | 12707 | Parietal_Sup_R | 26.0% Right_Superior_Parietal_Lobule; 19.0% Right_Lateral_Occipital_Cortex_superior_division |
| **3** | 38 | -29 | 37 | -7,30 | 12707 | Postcentral_R | 20.0% Right_Postcentral_Gyrus; 19.0% Right_Supramarginal_Gyrus_anterior_division |
| **4** | 35 | -53 | -28 | 16,34 | 11860 | Cerebelum_6_R | 0% no_label |
| **4** | 19 | -82 | -18 | 8,42 | 11860 | Cerebelum_6_R | 36.0% Right_Occipital_Fusiform_Gyrus; 7.0% Right_Lingual_Gyrus |
| **5** | -15 | -94 | -1 | 9,48 | 7050 | Calcarine_L | 35.0% Left_Occipital_Pole |
| **5** | 7 | -65 | 11 | 8,98 | 7050 | Calcarine_R | 68.0% Right_Intracalcarine_Cortex; 9.0% Right_Supracalcarine_Cortex; 6.0% Right_Lingual_Gyrus |
| **5** | 14 | -91 | 1 | 8,37 | 7050 | Calcarine_R | 48.0% Right_Occipital_Pole; 12.0% Right_Intracalcarine_Cortex; 7.0% Right_Lingual_Gyrus |
| **6** | -36 | -58 | -25 | 12,18 | 6435 | Cerebelum_6_L | 0% no_label |
| **6** | -34 | -82 | -23 | 7,72 | 6435 | Cerebelum_Crus1_L | 8.0% Left_Lateral_Occipital_Cortex_inferior_division; 6.0% Left_Occipital_Fusiform_Gyrus |
| **7** | -12 | -72 | 61 | 12,04 | 4878 | Precuneus_L | 63.0% Left_Lateral_Occipital_Cortex_superior_division |
| **7** | -5 | -51 | 73 | 8,42 | 4878 | Precuneus_L | 13.0% Left_Precuneous_Cortex; 8.0% Left_Postcentral_Gyrus; 6.0% Left_Superior_Parietal_Lobule |
| **8** | 42 | -60 | 16 | -10,12 | 4741 | Temporal_Mid_R | 22.0% Right_Lateral_Occipital_Cortex_superior_division; 21.0% Right_Angular_Gyrus; 20.0% Right_Lateral_Occipital_Cortex_inferior_division; 5.0% Right_Middle_Temporal_Gyrus_temporooccipital_part |
| **8** | 47 | -63 | -13 | -6,69 | 4741 | Occipital_Inf_R | 45.0% Right_Lateral_Occipital_Cortex_inferior_division; 18.0% Right_Inferior_Temporal_Gyrus_temporooccipital_part; 7.0% Right_Occipital_Fusiform_Gyrus; 6.0% Right_Temporal_Occipital_Fusiform_Cortex |
| **9** | -39 | -10 | 13 | -8,88 | 4290 | Insula_L | 52.0% Left_Central_Opercular_Cortex; 29.0% Left_Insular_Cortex |
| **9** | -53 | -34 | 16 | -7,84 | 4290 | Temporal_Sup_L | 51.0% Left_Planum_Temporale; 37.0% Left_Parietal_Operculum_Cortex |
| **10** | 45 | 18 | -13 | -8,50 | 4113 | OFCpost_R | 23.0% Right_Frontal_Orbital_Cortex; 19.0% Right_Temporal_Pole; 5.0% Right_Insular_Cortex |
| **10** | 30 | 45 | -13 | -8,39 | 4113 | OFCant_R | 78.0% Right_Frontal_Pole |
| **11** | -51 | -72 | 1 | -11,27 | 3894 | Occipital_Mid_L | 91.0% Left_Lateral_Occipital_Cortex_inferior_division |
| **12** | 40 | 54 | 6 | -9,87 | 3771 | Frontal_Mid_2_R | 91.0% Right_Frontal_Pole |
| **12** | 54 | 35 | 18 | -7,20 | 3771 | Frontal_Inf_Tri_R | 27.0% Right_Frontal_Pole; 19.0% Right_Inferior_Frontal_Gyrus_pars_triangularis; 9.0% Right_Middle_Frontal_Gyrus |
| **13** | 21 | -80 | 40 | -9,08 | 3498 | Occipital_Sup_R | 46.0% Right_Lateral_Occipital_Cortex_superior_division |
| **14** | -53 | 30 | 25 | 9,42 | 2528 | Frontal_Inf_Tri_L | 19.0% Left_Middle_Frontal_Gyrus; 9.0% Left_Inferior_Frontal_Gyrus_pars_triangularis |

| | | | | | | | |
|---|---|---|---|---|---|---|---|
| **14** | -36 | 40 | 42 | 5,74 | 2528 | Frontal_Mid_2_L | 11.0% Left_Frontal_Pole; 5.0% Left_Middle_Frontal_Gyrus |
| **15** | -65 | -51 | 1 | -10,31 | 2446 | Temporal_Mid_L | 74.0% Left_Middle_Temporal_Gyrus_temporooccipital_part; 12.0% Left_Angular_Gyrus |
| **16** | -58 | -41 | 32 | 13,77 | 2432 | SupraMarginal_L | 39.0% Left_Supramarginal_Gyrus_posterior_division; 34.0% Left_Supramarginal_Gyrus_anterior_division; 7.0% Left_Planum_Temporale |
| **17** | 35 | 9 | 28 | -10,94 | 2418 | Frontal_Inf_Oper_R | 15.0% Right_Precentral_Gyrus; 14.0% Right_Middle_Frontal_Gyrus; 12.0% Right_Inferior_Frontal_Gyrus_pars_opercularis |
| **17** | 59 | 16 | 18 | -7,22 | 2418 | Frontal_Inf_Oper_R | 68.0% Right_Inferior_Frontal_Gyrus_pars_opercularis; 11.0% Right_Precentral_Gyrus |
| **18** | 47 | -10 | 49 | -10,33 | 2391 | Precentral_R | 53.0% Right_Precentral_Gyrus; 17.0% Right_Postcentral_Gyrus |
| **19** | 2 | 30 | 52 | -8,51 | 1954 | Frontal_Sup_Medial_L | 63.0% Right_Superior_Frontal_Gyrus |
| **20** | 2 | -27 | 49 | -9,23 | 1722 | Cingulate_Mid_R | 40.0% Right_Precentral_Gyrus; 24.0% Right_Cingulate_Gyrus_posterior_division |
| **21** | 50 | -25 | -1 | 10,60 | 1708 | Temporal_Sup_R | 34.0% Right_Superior_Temporal_Gyrus_posterior_division; 9.0% Right_Middle_Temporal_Gyrus_posterior_division |
| **22** | 28 | 25 | 56 | -10,41 | 1694 | Frontal_Sup_2_R | 37.0% Right_Superior_Frontal_Gyrus; 24.0% Right_Middle_Frontal_Gyrus |
| **23** | -63 | -3 | 28 | -9,07 | 1640 | Postcentral_L | 24.0% Left_Postcentral_Gyrus; 23.0% Left_Precentral_Gyrus |
| **24** | -53 | 23 | 35 | 9,58 | 1640 | Frontal_Inf_Oper_L | 26.0% Left_Middle_Frontal_Gyrus |
| **25** | -22 | 9 | 4 | 8,18 | 1489 | Putamen_L | 98.0% Left_Putamen |
| **26** | -41 | -44 | 66 | -7,66 | 1366 | no_label | 16.0% Left_Superior_Parietal_Lobule; 5.0% Left_Postcentral_Gyrus |
| **27** | -15 | -84 | 40 | -6,68 | 1312 | Occipital_Sup_L | 48.0% Left_Lateral_Occipital_Cortex_superior_division |
| **28** | -20 | -6 | -16 | -8,61 | 1230 | Amygdala_L | 99.0% Left_Amygdala |
| **29** | -46 | 35 | -8 | -6,80 | 1093 | Frontal_Inf_Orb_2_L | 29.0% Left_Frontal_Orbital_Cortex; 13.0% Left_Inferior_Frontal_Gyrus_pars_triangularis; 10.0% Left_Frontal_Pole |
| **30** | 35 | 45 | 35 | 9,99 | 1025 | Frontal_Mid_2_R | 78.0% Right_Frontal_Pole |
| **31** | -58 | -27 | 52 | -9,68 | 984 | no_label | 19.0% Left_Postcentral_Gyrus; 12.0% Left_Supramarginal_Gyrus_anterior_division |
| **32** | 19 | -1 | -18 | -8,40 | 970 | Amygdala_R | 88.0% Right_Amygdala; 7.0% Right_Parahippocampal_Gyrus_anterior_division |
| **33** | -29 | -63 | 61 | -7,72 | 929 | Parietal_Sup_L | 55.0% Left_Lateral_Occipital_Cortex_superior_division; 8.0% Left_Superior_Parietal_Lobule |
| **34** | -29 | 61 | 4 | 9,02 | 874 | Frontal_Sup_2_L | 85.0% Left_Frontal_Pole |
| **35** | -41 | -60 | 30 | -7,81 | 833 | Angular_L | 27.0% Left_Angular_Gyrus; 17.0% Left_Lateral_Occipital_Cortex_superior_division |
| **36** | 45 | 25 | 6 | -8,05 | 820 | Frontal_Inf_Tri_R | 24.0% Right_Inferior_Frontal_Gyrus_pars_triangularis; 17.0% Right_Frontal_Operculum_Cortex; 6.0% Right_Inferior_Frontal_Gyrus_pars_opercularis |
| **37** | -8 | -48 | 1 | 7,08 | 779 | Lingual_L | 41.0% Left_Cingulate_Gyrus_posterior_division; 17.0% Left_Lingual_Gyrus |
| **38** | -44 | -56 | -13 | -6,83 | 779 | Fusiform_L | 39.0% Left_Inferior_Temporal_Gyrus_temporooccipital_part; 31.0% Left_Temporal_Occipital_Fusiform_Cortex; 6.0% Left_Lateral_Occipital_Cortex_inferior_division |
| **39** | 42 | -48 | -13 | -9,21 | 724 | Fusiform_R | 42.0% Right_Temporal_Occipital_Fusiform_Cortex; 16.0% Right_Inferior_Temporal_Gyrus_temporooccipital_part |
| **40** | -8 | -60 | 40 | 9,33 | 710 | Precuneus_L | 59.0% Left_Precuneous_Cortex |
| **41** | -36 | -44 | 37 | 9,21 | 615 | Parietal_Inf_L | 22.0% Left_Supramarginal_Gyrus_posterior_division; 16.0% Left_Superior_Parietal_Lobule; 8.0% Left_Supramarginal_Gyrus_anterior_division |
| **42** | -5 | 49 | 20 | -7,34 | 615 | Frontal_Sup_Medial_L | 67.0% Left_Paracingulate_Gyrus; 12.0% Left_Superior_Frontal_Gyrus |
| **43** | -55 | -68 | 13 | 11,78 | 601 | Temporal_Mid_L | 52.0% Left_Lateral_Occipital_Cortex_inferior_division; 34.0% Left_Lateral_Occipital_Cortex_superior_division |
| **44** | 52 | -10 | -28 | -7,66 | 588 | Temporal_Inf_R | 20.0% Right_Middle_Temporal_Gyrus_posterior_division; 12.0% Right_Inferior_Temporal_Gyrus_posterior_division; 12.0% Right_Inferior_Temporal_Gyrus_anterior_division; 6.0% Right_Middle_Temporal_Gyrus_anterior_division |
| **45** | -27 | -91 | -6 | -7,99 | 574 | Occipital_Inf_L | 27.0% Left_Occipital_Pole; 15.0% Left_Lateral_Occipital_Cortex_inferior_division; 9.0% Left_Occipital_Fusiform_Gyrus |

| | | | | | | | |
|---|---|---|---|---|---|---|---|
| **46** | 45 | 25 | 23 | -7,23 | 574 | Frontal_Inf_Tri_R | 34.0% Right_Middle_Frontal_Gyrus; 18.0% Right_Inferior_Frontal_Gyrus_pars_triangularis; 10.0% Right_Inferior_Frontal_Gyrus_pars_opercularis |
| **47** | -27 | 56 | 30 | -7,24 | 560 | Frontal_Sup_2_L | 48.0% Left_Frontal_Pole |
| **48** | -53 | 9 | 8 | 6,80 | 560 | Frontal_Inf_Oper_L | 42.0% Left_Inferior_Frontal_Gyrus_pars_opercularis; 24.0% Left_Precentral_Gyrus |
| **49** | 64 | -53 | 8 | -6,77 | 547 | Temporal_Mid_R | 68.0% Right_Middle_Temporal_Gyrus_temporooccipital_part |
| **50** | -60 | -53 | 40 | -6,64 | 492 | no_label | 27.0% Left_Supramarginal_Gyrus_posterior_division; 24.0% Left_Angular_Gyrus; 5.0% Left_Lateral_Occipital_Cortex_superior_division |
| **51** | 26 | 59 | 30 | -6,49 | 437 | Frontal_Sup_2_R | 40.0% Right_Frontal_Pole |
| **52** | -32 | 28 | 4 | 9,24 | 383 | Insula_L | 19.0% Left_Frontal_Orbital_Cortex; 19.0% Left_Insular_Cortex; 17.0% Left_Inferior_Frontal_Gyrus_pars_triangularis; 13.0% Left_Frontal_Operculum_Cortex |
| **53** | 64 | -29 | -18 | -5,96 | 369 | Temporal_Inf_R | 35.0% Right_Inferior_Temporal_Gyrus_posterior_division; 34.0% Right_Middle_Temporal_Gyrus_posterior_division |
| **54** | -39 | -15 | 44 | -8,50 | 369 | Postcentral_L | 45.0% Left_Precentral_Gyrus; 17.0% Left_Postcentral_Gyrus |
| **55** | 50 | -39 | 42 | -6,24 | 355 | SupraMarginal_R | 46.0% Right_Supramarginal_Gyrus_posterior_division; 7.0% Right_Angular_Gyrus |
| **56** | 30 | 49 | 25 | -7,52 | 342 | Frontal_Mid_2_R | 85.0% Right_Frontal_Pole |
| **57** | 14 | -60 | 61 | 7,23 | 342 | Precuneus_R | 43.0% Right_Lateral_Occipital_Cortex_superior_division; 13.0% Right_Superior_Parietal_Lobule; 8.0% Right_Precuneous_Cortex |
| **58** | -5 | -1 | 4 | 7,40 | 328 | no_label | 56.0% Left_Thalamus; 28.0% Left_Lateral_Ventrical |
| **59** | -1 | -37 | -37 | 6,48 | 314 | no_label | 99.0% Brain-Stem |
| **60** | -55 | -53 | 16 | -7,11 | 314 | Temporal_Mid_L | 41.0% Left_Angular_Gyrus; 22.0% Left_Supramarginal_Gyrus_posterior_division |
| **61** | -36 | 11 | 8 | 7,43 | 314 | Insula_L | 48.0% Left_Frontal_Operculum_Cortex; 24.0% Left_Central_Opercular_Cortex; 14.0% Left_Insular_Cortex |
| **62** | 2 | -44 | 8 | 7,81 | 314 | no_label | 19.0% Right_Cingulate_Gyrus_posterior_division |
| **63** | 28 | -15 | 56 | 6,61 | 301 | Precentral_R | 47.0% Right_Precentral_Gyrus |
| **64** | 14 | -48 | 37 | -7,62 | 301 | Precuneus_R | 19.0% Right_Precuneous_Cortex; 18.0% Right_Cingulate_Gyrus_posterior_division |
| **65** | 23 | 42 | 40 | 7,08 | 287 | Frontal_Sup_2_R | 65.0% Right_Frontal_Pole; 5.0% Right_Superior_Frontal_Gyrus |
| **66** | -22 | -68 | -28 | 6,80 | 287 | Cerebelum_6_L | 0% no_label |
| **67** | 28 | -48 | -6 | -6,03 | 287 | Lingual_R | 63.0% Right_Lingual_Gyrus; 25.0% Right_Temporal_Occipital_Fusiform_Cortex |
| **68** | 16 | -3 | -4 | 7,86 | 287 | no_label | 94.0% Right_Pallidum |
| **69** | -15 | -58 | 18 | 6,39 | 273 | Cuneus_L | 54.0% Left_Precuneous_Cortex; 11.0% Left_Supracalcarine_Cortex |
| **70** | -29 | 18 | -13 | -7,23 | 260 | Insula_L | 56.0% Left_Frontal_Orbital_Cortex; 23.0% Left_Insular_Cortex |
| **71** | -27 | -70 | 42 | 8,94 | 246 | Parietal_Inf_L | 66.0% Left_Lateral_Occipital_Cortex_superior_division |
| **72** | -5 | -75 | -16 | 7,12 | 232 | Cerebelum_6_L | 7.0% Left_Lingual_Gyrus |
| **73** | -55 | -46 | 52 | -7,58 | 232 | no_label | 53.0% Left_Supramarginal_Gyrus_posterior_division; 8.0% Left_Angular_Gyrus; 6.0% Left_Supramarginal_Gyrus_anterior_division |
| **74** | -29 | -34 | 76 | -6,96 | 205 | Postcentral_L | 10.0% Left_Postcentral_Gyrus |
| **75** | -10 | 56 | -16 | -6,11 | 191 | OFCmed_L | 23.0% Left_Frontal_Pole |
| **76** | -29 | -75 | 25 | -6,68 | 191 | Occipital_Mid_L | 66.0% Left_Lateral_Occipital_Cortex_superior_division |
| **77** | 19 | 33 | -20 | -6,35 | 178 | OFCmed_R | 47.0% Right_Frontal_Orbital_Cortex; 38.0% Right_Frontal_Pole |
| **78** | 9 | 42 | 54 | -6,49 | 164 | Frontal_Sup_Medial_R | 51.0% Right_Frontal_Pole; 18.0% Right_Superior_Frontal_Gyrus |
| **79** | 4 | 42 | 28 | -6,14 | 164 | Cingulate_Ant_R | 90.0% Right_Paracingulate_Gyrus |
| **80** | 4 | -48 | -4 | 6,35 | 164 | Vermis_4_5 | 0% no_label |
| **81** | -55 | -41 | 11 | 7,33 | 164 | Temporal_Mid_L | 18.0% Left_Supramarginal_Gyrus_posterior_division; 13.0% Left_Superior_Temporal_Gyrus_posterior_division |
| **82** | 52 | -63 | 42 | -7,27 | 150 | Angular_R | 68.0% Right_Lateral_Occipital_Cortex_superior_division; 11.0% Right_Angular_Gyrus |
| **83** | 4 | -65 | -1 | 5,70 | 137 | Lingual_R | 54.0% Right_Lingual_Gyrus |
| **84** | -55 | -8 | 47 | 8,00 | 137 | Postcentral_L | 65.0% Left_Precentral_Gyrus; 12.0% Left_Postcentral_Gyrus |

| | | | | | | | |
|---|---|---|---|---|---|---|---|
| **85** | -8 | 21 | 61 | -6,22 | 137 | Supp_Motor_Area_L | 43.0% Left_Superior_Frontal_Gyrus |
| **86** | -41 | -20 | 4 | -7,55 | 137 | Temporal_Sup_L | 63.0% Left_Heschl's_Gyrus_(includes_H1_and_H2); 8.0% Left_Planum_Polare |
| **87** | -15 | -15 | 8 | 6,27 | 123 | Thalamus_L | 99.0% Left_Thalamus |
| **88** | -8 | -41 | 54 | -6,17 | 123 | Cingulate_Mid_L | 33.0% Left_Postcentral_Gyrus; 29.0% Left_Precuneous_Cortex; 6.0% Left_Precentral_Gyrus |
| **89** | -51 | -10 | 8 | -6,23 | 123 | Heschl_L | 56.0% Left_Central_Opercular_Cortex; 17.0% Left_Heschl's_Gyrus_(includes_H1_and_H2); 13.0% Left_Planum_Polare |
| **90** | 40 | -3 | 56 | 5,66 | 123 | Frontal_Mid_2_R | 39.0% Right_Precentral_Gyrus; 21.0% Right_Middle_Frontal_Gyrus |
| **91** | 2 | -32 | -4 | 5,82 | 123 | no_label | 89.0% Brain-Stem |
| **92** | -1 | -10 | 47 | 6,26 | 123 | Cingulate_Mid_L | 41.0% Left_Juxtapositional_Lobule_Cortex_(formerly_Supplementary_Motor_Cortex); 28.0% Left_Cingulate_Gyrus_anterior_division; 6.0% Left_Precentral_Gyrus |
| **93** | -24 | -58 | 1 | 7,12 | 123 | no_label | 14.0% Left_Lingual_Gyrus; 6.0% Left_Precuneous_Cortex |
| **94** | 9 | 18 | 37 | 6,07 | 123 | Cingulate_Mid_R | 47.0% Right_Paracingulate_Gyrus; 32.0% Right_Cingulate_Gyrus_anterior_division |
| **95** | -10 | 28 | -16 | -6,18 | 109 | Rectus_L | 8.0% Left_Subcallosal_Cortex |
| **96** | -10 | -34 | 76 | 5,71 | 109 | Paracentral_Lobule_L | 39.0% Left_Postcentral_Gyrus; 20.0% Left_Precentral_Gyrus |
| **97** | 14 | -41 | 56 | 6,94 | 109 | Precuneus_R | 31.0% Right_Postcentral_Gyrus; 6.0% Right_Precuneous_Cortex |
| **98** | -46 | 49 | 6 | -5,77 | 109 | Frontal_Mid_2_L | 81.0% Left_Frontal_Pole |
| **99** | -1 | 18 | -25 | -5,97 | 109 | no_label | 17.0% Left_Subcallosal_Cortex |
| **100** | 38 | -70 | -13 | -6,02 | 109 | Fusiform_R | 33.0% Right_Occipital_Fusiform_Gyrus; 22.0% Right_Lateral_Occipital_Cortex_inferior_division |
| **101** | -60 | -15 | -30 | -6,19 | 109 | no_label | 36.0% Left_Inferior_Temporal_Gyrus_posterior_division; 26.0% Left_Middle_Temporal_Gyrus_posterior_division; 17.0% Left_Inferior_Temporal_Gyrus_anterior_division; 6.0% Left_Middle_Temporal_Gyrus_anterior_division |
| **102** | -8 | -103 | 4 | 6,38 | 96 | Occipital_Sup_L | 59.0% Left_Occipital_Pole |
| **103** | -8 | -46 | 78 | 7,28 | 96 | Precuneus_L | 29.0% Left_Postcentral_Gyrus |
| **104** | 69 | -37 | -4 | -5,90 | 96 | Temporal_Mid_R | 49.0% Right_Middle_Temporal_Gyrus_posterior_division; 33.0% Right_Middle_Temporal_Gyrus_temporooccipital_part |
| **105** | -3 | -94 | -6 | 5,43 | 96 | Calcarine_L | 48.0% Left_Occipital_Pole; 9.0% Left_Lingual_Gyrus; 6.0% Left_Intracalcarine_Cortex |
| **106** | 23 | -1 | 56 | 6,59 | 96 | Frontal_Sup_2_R | 40.0% Right_Superior_Frontal_Gyrus; 12.0% Right_Middle_Frontal_Gyrus; 10.0% Right_Precentral_Gyrus |
| **107** | -53 | -10 | -13 | -5,77 | 96 | Temporal_Mid_L | 19.0% Left_Middle_Temporal_Gyrus_anterior_division; 19.0% Left_Superior_Temporal_Gyrus_anterior_division; 17.0% Left_Middle_Temporal_Gyrus_posterior_division; 15.0% Left_Superior_Temporal_Gyrus_posterior_division |
| **108** | 45 | 2 | 54 | 6,45 | 96 | Frontal_Mid_2_R | 45.0% Right_Middle_Frontal_Gyrus; 25.0% Right_Precentral_Gyrus |
| **109** | -55 | -8 | -28 | -5,61 | 96 | Temporal_Inf_L | 39.0% Left_Middle_Temporal_Gyrus_anterior_division; 21.0% Left_Middle_Temporal_Gyrus_posterior_division; 13.0% Left_Inferior_Temporal_Gyrus_anterior_division; 6.0% Left_Inferior_Temporal_Gyrus_posterior_division |
| **110** | -1 | -53 | -32 | 5,63 | 82 | Vermis_9 | 0% no_label |
| **111** | 52 | 35 | -8 | -5,98 | 82 | Frontal_Inf_Orb_2_R | 48.0% Right_Frontal_Pole; 19.0% Right_Inferior_Frontal_Gyrus_pars_triangularis; 13.0% Right_Frontal_Orbital_Cortex |
| **112** | 54 | 33 | 1 | 6,14 | 82 | Frontal_Inf_Tri_R | 51.0% Right_Inferior_Frontal_Gyrus_pars_triangularis; 31.0% Right_Frontal_Pole |
| **113** | 50 | -22 | 40 | -5,72 | 82 | Postcentral_R | 47.0% Right_Postcentral_Gyrus; 22.0% Right_Supramarginal_Gyrus_anterior_division |
| **114** | -55 | -15 | -16 | -5,83 | 82 | Temporal_Mid_L | 31.0% Left_Middle_Temporal_Gyrus_posterior_division; 12.0% Left_Middle_Temporal_Gyrus_anterior_division; 5.0% Left_Superior_Temporal_Gyrus_posterior_division |
| **115** | 59 | -56 | 40 | -6,35 | 82 | Parietal_Inf_R | 38.0% Right_Angular_Gyrus; 15.0% Right_Lateral_Occipital_Cortex_superior_division |
| **116** | 42 | 2 | 4 | -5,36 | 82 | Insula_R | 66.0% Right_Insular_Cortex; 15.0% Right_Central_Opercular_Cortex |

| | | | | | | | |
|---|---|---|---|---|---|---|---|
| **117** | -1 | -44 | -13 | 5,98 | 82 | Vermis_3 | 0% no_label |
| **118** | -48 | -32 | 54 | -5,77 | 82 | Postcentral_L | 41.0% Left_Postcentral_Gyrus; 14.0% Left_Superior_Parietal_Lobule; 11.0% Left_Supramarginal_Gyrus_anterior_division |
| **119** | -58 | 14 | -1 | 6,59 | 82 | no_label | 14.0% Left_Inferior_Frontal_Gyrus_pars_opercularis |
| **120** | 11 | 16 | 56 | -5,84 | 82 | Supp_Motor_Area_R | 13.0% Right_Superior_Frontal_Gyrus |
| **121** | -5 | 40 | -20 | -5,47 | 68 | Rectus_L | 47.0% Left_Frontal_Medial_Cortex |
| **122** | 42 | -8 | -8 | -5,72 | 68 | Insula_R | 47.0% Right_Insular_Cortex; 27.0% Right_Planum_Polare |
| **123** | -58 | 11 | 35 | 5,83 | 68 | Precentral_L | 16.0% Left_Precentral_Gyrus |
| **124** | 33 | -70 | 49 | -5,93 | 68 | Parietal_Sup_R | 62.0% Right_Lateral_Occipital_Cortex_superior_division |
| **125** | -1 | 30 | 13 | -6,60 | 68 | Cingulate_Ant_L | 71.0% Left_Cingulate_Gyrus_anterior_division |
| **126** | -10 | -44 | 64 | -5,97 | 68 | Precuneus_L | 44.0% Left_Postcentral_Gyrus; 16.0% Left_Precuneous_Cortex |
| **127** | -5 | 21 | -23 | -5,59 | 68 | Rectus_L | 64.0% Left_Subcallosal_Cortex |
| **128** | -10 | -17 | 52 | -6,28 | 68 | Supp_Motor_Area_L | 14.0% Left_Precentral_Gyrus; 9.0% Left_Juxtapositional_Lobule_Cortex_(formerly_Supplementary_Motor_Cortex) |
| **129** | 28 | -94 | 11 | 5,99 | 68 | Occipital_Mid_R | 55.0% Right_Occipital_Pole; 9.0% Right_Lateral_Occipital_Cortex_superior_division |
| **130** | -36 | -91 | 16 | -6,14 | 68 | no_label | 39.0% Left_Occipital_Pole; 25.0% Left_Lateral_Occipital_Cortex_superior_division |
| **131** | -29 | 35 | 49 | 7,61 | 68 | Frontal_Mid_2_L | 24.0% Left_Middle_Frontal_Gyrus; 8.0% Left_Frontal_Pole |
| **132** | -8 | -22 | 42 | 5,64 | 68 | Cingulate_Mid_L | 63.0% Left_Cingulate_Gyrus_posterior_division; 14.0% Left_Precentral_Gyrus |
| **133** | -44 | 47 | -11 | -5,93 | 68 | Frontal_Inf_Orb_2_L | 86.0% Left_Frontal_Pole |
| **134** | -53 | -25 | -11 | -6,13 | 68 | Temporal_Mid_L | 47.0% Left_Middle_Temporal_Gyrus_posterior_division |
| **135** | -67 | -39 | 16 | -5,86 | 68 | no_label | 17.0% Left_Superior_Temporal_Gyrus_posterior_division; 14.0% Left_Supramarginal_Gyrus_posterior_division; 6.0% Left_Planum_Temporale |
| **136** | 2 | -75 | 37 | 6,79 | 68 | Precuneus_L | 41.0% Right_Precuneous_Cortex; 17.0% Right_Cuneal_Cortex; 13.0% Left_Precuneous_Cortex; 6.0% Left_Cuneal_Cortex |
| **137** | -41 | 18 | -23 | -5,78 | 68 | Temporal_Pole_Sup_L | 72.0% Left_Temporal_Pole; 10.0% Left_Frontal_Orbital_Cortex |

***Table S3.*** *Regions whose activation changed positively or negatively over time when regulating to ECN. X-, Y- and Z-coordinates are in MNI space; volume is in mm.*

| # | X | Y | Z | Z-STAT | VOLUME | AAL | HARVARD-OXFORD |
|---|---|---|---|---|---|---|---|
| **1** | 45 | -37 | 59 | -13,56 | 66799 | Postcentral_R | 21.0% Right_Superior_Parietal_Lobule; 21.0% Right_Postcentral_Gyrus; 8.0% Right_Supramarginal_Gyrus_posterior_division |
| **1** | 59 | 16 | 18 | -11,42 | 66799 | Frontal_Inf_Oper_R | 68.0% Right_Inferior_Frontal_Gyrus_pars_opercularis; 11.0% Right_Precentral_Gyrus |
| **1** | 40 | 9 | 4 | -11,41 | 66799 | Insula_R | 36.0% Right_Central_Opercular_Cortex; 26.0% Right_Insular_Cortex; 15.0% Right_Frontal_Operculum_Cortex |
| **1** | 59 | -29 | 20 | -11,20 | 66799 | Temporal_Sup_R | 44.0% Right_Parietal_Operculum_Cortex; 26.0% Right_Planum_Temporale; 10.0% Right_Supramarginal_Gyrus_anterior_division |
| **1** | 28 | -53 | 52 | -11,00 | 66799 | Parietal_Inf_R | 38.0% Right_Superior_Parietal_Lobule |
| **1** | 35 | 9 | 30 | -10,73 | 66799 | Frontal_Inf_Oper_R | 22.0% Right_Middle_Frontal_Gyrus; 21.0% Right_Precentral_Gyrus; 14.0% Right_Inferior_Frontal_Gyrus_pars_opercularis |
| **1** | 50 | -10 | 49 | -9,92 | 66799 | Precentral_R | 60.0% Right_Precentral_Gyrus; 18.0% Right_Postcentral_Gyrus |
| **1** | 64 | -29 | 44 | -9,76 | 66799 | SupraMarginal_R | 57.0% Right_Supramarginal_Gyrus_anterior_division |
| **1** | 40 | -13 | 20 | -8,77 | 66799 | Rolandic_Oper_R | 73.0% Right_Central_Opercular_Cortex; 9.0% Right_Parietal_Operculum_Cortex |
| **1** | 21 | -20 | 76 | -8,57 | 66799 | Precentral_R | 46.0% Right_Precentral_Gyrus |
| **1** | 64 | -8 | 8 | -7,32 | 66799 | Temporal_Sup_R | 43.0% Right_Central_Opercular_Cortex; 13.0% Right_Planum_Temporale; 9.0% Right_Planum_Polare; 7.0% Right_Postcentral_Gyrus; 6.0% Right_Superior_Temporal_Gyrus_anterior_division |
| **1** | 45 | 9 | -20 | -5,79 | 66799 | Temporal_Pole_Sup_R | 67.0% Right_Temporal_Pole |
| **2** | -67 | -27 | 37 | -12,04 | 43941 | no_label | 15.0% Left_Supramarginal_Gyrus_anterior_division |
| **2** | -46 | 6 | -1 | -12,01 | 43941 | Insula_L | 53.0% Left_Central_Opercular_Cortex; 12.0% Left_Frontal_Operculum_Cortex; 8.0% Left_Insular_Cortex |
| **2** | -53 | -34 | 16 | -11,50 | 43941 | Temporal_Sup_L | 51.0% Left_Planum_Temporale; 37.0% Left_Parietal_Operculum_Cortex |
| **2** | -53 | -37 | 56 | -10,06 | 43941 | no_label | 23.0% Left_Supramarginal_Gyrus_anterior_division; 8.0% Left_Supramarginal_Gyrus_posterior_division; 6.0% Left_Postcentral_Gyrus |
| **2** | -53 | 37 | 1 | -9,78 | 43941 | Frontal_Inf_Tri_L | 40.0% Left_Frontal_Pole; 35.0% Left_Inferior_Frontal_Gyrus_pars_triangularis |
| **2** | -39 | -15 | 44 | -9,55 | 43941 | Postcentral_L | 45.0% Left_Precentral_Gyrus; 17.0% Left_Postcentral_Gyrus |
| **2** | -24 | -48 | 64 | -9,33 | 43941 | Parietal_Sup_L | 50.0% Left_Superior_Parietal_Lobule; 10.0% Left_Postcentral_Gyrus |
| **2** | -29 | -13 | 11 | -7,32 | 43941 | no_label | 50.0% Left_Putamen |
| **2** | -63 | -6 | 13 | -7,31 | 43941 | Postcentral_L | 39.0% Left_Postcentral_Gyrus; 31.0% Left_Precentral_Gyrus |
| **2** | -24 | -25 | 76 | -6,38 | 43941 | Precentral_L | 30.0% Left_Precentral_Gyrus; 6.0% Left_Postcentral_Gyrus |
| **3** | 2 | -15 | 59 | -10,87 | 19798 | Supp_Motor_Area_R | 43.0% Right_Precentral_Gyrus; 23.0% Right_Juxtapositional_Lobule_Cortex_(formerly_Supplementary_Motor_Cortex) |
| **3** | 2 | 16 | 56 | -9,75 | 19798 | Supp_Motor_Area_R | 45.0% Right_Superior_Frontal_Gyrus; 10.0% Right_Paracingulate_Gyrus; 6.0% Left_Superior_Frontal_Gyrus |
| **3** | -1 | 23 | 23 | -8,37 | 19798 | Cingulate_Ant_L | 92.0% Left_Cingulate_Gyrus_anterior_division |
| **3** | -12 | -34 | 47 | -7,80 | 19798 | Cingulate_Mid_L | 38.0% Left_Precentral_Gyrus; 15.0% Left_Cingulate_Gyrus_posterior_division; 11.0% Left_Postcentral_Gyrus; 10.0% Left_Precuneous_Cortex |
| **4** | 2 | -60 | 32 | 10,82 | 8061 | Precuneus_L | 91.0% Right_Precuneous_Cortex |
| **4** | -17 | -58 | 18 | 9,66 | 8061 | Cuneus_L | 43.0% Left_Precuneous_Cortex; 16.0% Left_Supracalcarine_Cortex |
| **5** | 50 | -51 | -20 | -9,78 | 7228 | Temporal_Inf_R | 75.0% Right_Inferior_Temporal_Gyrus_temporooccipital_part; 13.0% Right_Temporal_Occipital_Fusiform_Cortex |
| **5** | 47 | -65 | 1 | -9,54 | 7228 | Temporal_Mid_R | 62.0% Right_Lateral_Occipital_Cortex_inferior_division; 8.0% Right_Middle_Temporal_Gyrus_temporooccipital_part |
| **6** | -46 | -70 | -1 | -10,72 | 4973 | Occipital_Mid_L | 66.0% Left_Lateral_Occipital_Cortex_inferior_division |
| **7** | -65 | -46 | 6 | -10,37 | 4727 | Temporal_Mid_L | 43.0% Left_Middle_Temporal_Gyrus_temporooccipital_part; 22.0% Left_Supramarginal_Gyrus_posterior_division; 11.0% Left_Angular_Gyrus; 7.0% Left_Middle_Temporal_Gyrus_posterior_division; 5.0% Left_Superior_Temporal_Gyrus_posterior_division |
| **8** | 19 | -80 | 40 | -10,30 | 4400 | Cuneus_R | 48.0% Right_Lateral_Occipital_Cortex_superior_division; 7.0% Right_Cuneal_Cortex |
| **9** | 47 | 45 | 20 | -9,58 | 3552 | Frontal_Mid_2_R | 72.0% Right_Frontal_Pole |

| | | | | | | | |
|---|---|---|---|---|---|---|---|
| **9** | 23 | 61 | 30 | -8,52 | 3552 | no_label | 40.0% Right_Frontal_Pole |
| **10** | -15 | -75 | 61 | 12,88 | 1913 | Precuneus_L | 25.0% Left_Lateral_Occipital_Cortex_superior_division |
| **11** | -17 | -75 | 35 | -7,69 | 1640 | Cuneus_L | 22.0% Left_Cuneal_Cortex; 18.0% Left_Lateral_Occipital_Cortex_superior_division; 17.0% Left_Precuneous_Cortex |
| **12** | -22 | -101 | -6 | -11,10 | 1571 | Occipital_Inf_L | 78.0% Left_Occipital_Pole |
| **13** | -27 | 56 | 30 | -8,78 | 1544 | Frontal_Sup_2_L | 48.0% Left_Frontal_Pole |
| **14** | -34 | -53 | -16 | -11,50 | 1298 | Fusiform_L | 62.0% Left_Temporal_Occipital_Fusiform_Cortex |
| **15** | 30 | -46 | -18 | -7,76 | 1216 | Fusiform_R | 88.0% Right_Temporal_Occipital_Fusiform_Cortex |
| **16** | -15 | -94 | -6 | 9,11 | 1148 | Calcarine_L | 46.0% Left_Occipital_Pole; 6.0% Left_Lateral_Occipital_Cortex_inferior_division |
| **17** | 19 | 35 | -23 | -7,28 | 1134 | OFCmed_R | 50.0% Right_Frontal_Pole; 25.0% Right_Frontal_Orbital_Cortex |
| **18** | -34 | -56 | 52 | -8,22 | 1134 | Parietal_Inf_L | 36.0% Left_Superior_Parietal_Lobule; 11.0% Left_Lateral_Occipital_Cortex_superior_division; 9.0% Left_Supramarginal_Gyrus_posterior_division; 8.0% Left_Angular_Gyrus |
| **19** | -51 | 6 | 42 | -9,21 | 1107 | Precentral_L | 38.0% Left_Middle_Frontal_Gyrus; 36.0% Left_Precentral_Gyrus |
| **20** | 26 | 2 | -13 | -7,76 | 1093 | Amygdala_R | 33.0% Right_Amygdala |
| **21** | -22 | 2 | 54 | 7,66 | 820 | Frontal_Sup_2_L | 42.0% Left_Superior_Frontal_Gyrus; 10.0% Left_Middle_Frontal_Gyrus |
| **22** | -12 | 45 | 47 | -8,80 | 724 | Frontal_Sup_2_L | 75.0% Left_Frontal_Pole; 8.0% Left_Superior_Frontal_Gyrus |
| **23** | -53 | -6 | -32 | -8,73 | 710 | Temporal_Inf_L | 31.0% Left_Inferior_Temporal_Gyrus_anterior_division; 29.0% Left_Middle_Temporal_Gyrus_anterior_division; 11.0% Left_Middle_Temporal_Gyrus_posterior_division; 7.0% Left_Inferior_Temporal_Gyrus_posterior_division |
| **24** | 19 | -87 | -6 | 7,97 | 697 | Lingual_R | 32.0% Right_Occipital_Fusiform_Gyrus; 16.0% Right_Occipital_Pole; 16.0% Right_Lingual_Gyrus |
| **25** | 4 | 49 | 35 | 7,32 | 683 | Frontal_Sup_Medial_R | 79.0% Right_Superior_Frontal_Gyrus; 5.0% Right_Paracingulate_Gyrus |
| **26** | 52 | -8 | -18 | 7,59 | 615 | Temporal_Mid_R | 27.0% Right_Middle_Temporal_Gyrus_posterior_division; 18.0% Right_Middle_Temporal_Gyrus_anterior_division; 11.0% Right_Superior_Temporal_Gyrus_anterior_division; 9.0% Right_Superior_Temporal_Gyrus_posterior_division |
| **27** | 33 | -91 | -4 | -6,99 | 574 | Occipital_Inf_R | 40.0% Right_Occipital_Pole; 29.0% Right_Lateral_Occipital_Cortex_inferior_division |
| **28** | -22 | 23 | -18 | -8,54 | 574 | OFCpost_L | 82.0% Left_Frontal_Orbital_Cortex |
| **29** | 50 | 49 | 1 | -8,53 | 506 | Frontal_Mid_2_R | 46.0% Right_Frontal_Pole |
| **30** | 2 | -10 | 73 | -8,19 | 492 | Supp_Motor_Area_R | 14.0% Right_Juxtapositional_Lobule_Cortex_(formerly_Supplementary_Motor_Cortex); 7.0% Right_Precentral_Gyrus |
| **31** | -53 | -80 | 11 | -7,42 | 342 | no_label | 18.0% Left_Lateral_Occipital_Cortex_inferior_division |
| **32** | 4 | -44 | 11 | 8,64 | 328 | Cingulate_Post_R | 49.0% Right_Cingulate_Gyrus_posterior_division |
| **33** | -3 | -75 | 54 | 7,31 | 328 | Precuneus_L | 59.0% Left_Precuneous_Cortex; 9.0% Left_Lateral_Occipital_Cortex_superior_division |
| **34** | -3 | -70 | -32 | -6,67 | 328 | Cerebelum_8_L | 0% no_label |
| **35** | -32 | -82 | 47 | 8,19 | 287 | no_label | 28.0% Left_Lateral_Occipital_Cortex_superior_division |
| **36** | 11 | -17 | 6 | -5,97 | 287 | Thalamus_R | 100.0% Right_Thalamus |
| **37** | -22 | 2 | -16 | -7,36 | 273 | Amygdala_L | 17.0% Left_Amygdala; 5.0% Left_Parahippocampal_Gyrus_anterior_division |
| **38** | -20 | -15 | 20 | -6,40 | 273 | no_label | 0% no_label |
| **39** | -1 | -34 | 35 | 6,53 | 260 | Cingulate_Mid_L | 91.0% Left_Cingulate_Gyrus_posterior_division |
| **40** | -24 | -101 | 6 | 7,25 | 246 | Occipital_Mid_L | 64.0% Left_Occipital_Pole |
| **41** | 38 | -70 | -16 | -7,35 | 232 | Fusiform_R | 52.0% Right_Occipital_Fusiform_Gyrus; 27.0% Right_Lateral_Occipital_Cortex_inferior_division |
| **42** | -3 | -58 | 61 | 6,88 | 232 | Precuneus_L | 78.0% Left_Precuneous_Cortex |
| **43** | 11 | 64 | 1 | 7,54 | 219 | Frontal_Sup_Medial_R | 32.0% Right_Frontal_Pole |
| **44** | -65 | -8 | -16 | 6,68 | 219 | Temporal_Mid_L | 44.0% Left_Middle_Temporal_Gyrus_anterior_division; 28.0% Left_Middle_Temporal_Gyrus_posterior_division; 5.0% Left_Superior_Temporal_Gyrus_posterior_division |

| | | | | | | | |
|---|---|---|---|---|---|---|---|
| **45** | 26 | 11 | 56 | -6,62 | 205 | Frontal_Sup_2_R | 39.0% Right_Superior_Frontal_Gyrus; 28.0% Right_Middle_Frontal_Gyrus |
| **46** | -41 | 2 | 56 | -6,86 | 205 | Precentral_L | 49.0% Left_Middle_Frontal_Gyrus; 23.0% Left_Precentral_Gyrus |
| **47** | -41 | -75 | 16 | -7,25 | 205 | Occipital_Mid_L | 36.0% Left_Lateral_Occipital_Cortex_superior_division; 29.0% Left_Lateral_Occipital_Cortex_inferior_division |
| **48** | -34 | -72 | -18 | -7,32 | 205 | Fusiform_L | 63.0% Left_Occipital_Fusiform_Gyrus; 12.0% Left_Lateral_Occipital_Cortex_inferior_division |
| **49** | 21 | 35 | 42 | 6,07 | 191 | Frontal_Sup_2_R | 35.0% Right_Frontal_Pole; 32.0% Right_Superior_Frontal_Gyrus; 13.0% Right_Middle_Frontal_Gyrus |
| **50** | -12 | 56 | -18 | -6,62 | 191 | OFCmed_L | 44.0% Left_Frontal_Pole |
| **51** | 19 | -82 | -18 | 8,00 | 178 | Cerebelum_6_R | 36.0% Right_Occipital_Fusiform_Gyrus; 7.0% Right_Lingual_Gyrus |
| **52** | 14 | -77 | -32 | -6,71 | 178 | Cerebelum_Crus1_R | 0% no_label |
| **53** | -27 | 18 | 47 | 6,03 | 178 | Frontal_Mid_2_L | 32.0% Left_Middle_Frontal_Gyrus; 26.0% Left_Superior_Frontal_Gyrus |
| **54** | -63 | -3 | 25 | -7,09 | 178 | Postcentral_L | 33.0% Left_Precentral_Gyrus; 29.0% Left_Postcentral_Gyrus |
| **55** | 64 | -51 | 23 | 7,26 | 164 | Temporal_Sup_R | 58.0% Right_Angular_Gyrus |
| **56** | 11 | -53 | 47 | 6,84 | 150 | Precuneus_R | 52.0% Right_Precuneous_Cortex |
| **57** | -27 | -58 | 8 | -6,53 | 150 | Calcarine_L | 47.0% Left_Lateral_Ventrical; 9.0% Left_Precuneous_Cortex; 6.0% Left_Intracalcarine_Cortex; 5.0% Left_Lingual_Gyrus |
| **58** | 47 | 2 | -35 | -5,68 | 150 | Temporal_Inf_R | 36.0% Right_Inferior_Temporal_Gyrus_anterior_division; 14.0% Right_Middle_Temporal_Gyrus_anterior_division; 11.0% Right_Temporal_Pole |
| **59** | -36 | 30 | -20 | -6,34 | 123 | OFCpost_L | 71.0% Left_Frontal_Orbital_Cortex; 11.0% Left_Frontal_Pole |
| **60** | -32 | 18 | 11 | -6,06 | 123 | Insula_L | 67.0% Left_Frontal_Operculum_Cortex; 10.0% Left_Insular_Cortex |
| **61** | 50 | -56 | 40 | 6,17 | 123 | Angular_R | 53.0% Right_Angular_Gyrus; 17.0% Right_Lateral_Occipital_Cortex_superior_division |
| **62** | -55 | -63 | 28 | -5,91 | 123 | Angular_L | 64.0% Left_Lateral_Occipital_Cortex_superior_division; 27.0% Left_Angular_Gyrus |
| **63** | -8 | -84 | -11 | 6,53 | 109 | Lingual_L | 39.0% Left_Lingual_Gyrus; 19.0% Left_Occipital_Fusiform_Gyrus |
| **64** | -8 | -48 | 1 | 7,51 | 109 | Lingual_L | 41.0% Left_Cingulate_Gyrus_posterior_division; 17.0% Left_Lingual_Gyrus |
| **65** | -39 | 21 | 28 | 6,94 | 109 | Frontal_Inf_Tri_L | 38.0% Left_Middle_Frontal_Gyrus; 6.0% Left_Inferior_Frontal_Gyrus_pars_triangularis; 5.0% Left_Inferior_Frontal_Gyrus_pars_opercularis |
| **66** | -34 | 11 | 35 | -5,82 | 109 | Frontal_Mid_2_L | 30.0% Left_Middle_Frontal_Gyrus |
| **67** | 35 | 14 | -37 | -7,32 | 109 | Temporal_Pole_Mid_R | 46.0% Right_Temporal_Pole |
| **68** | -46 | -60 | -20 | -5,60 | 109 | Fusiform_L | 41.0% Left_Temporal_Occipital_Fusiform_Cortex; 28.0% Left_Inferior_Temporal_Gyrus_temporooccipital_part; 12.0% Left_Lateral_Occipital_Cortex_inferior_division; 7.0% Left_Occipital_Fusiform_Gyrus |
| **69** | -3 | 16 | 4 | -6,01 | 109 | no_label | 78.0% Left_Lateral_Ventrical |
| **70** | 54 | 42 | -1 | -7,12 | 109 | Frontal_Inf_Tri_R | 36.0% Right_Frontal_Pole; 5.0% Right_Inferior_Frontal_Gyrus_pars_triangularis |
| **71** | -27 | -77 | 28 | -5,59 | 109 | Occipital_Mid_L | 79.0% Left_Lateral_Occipital_Cortex_superior_division |
| **72** | -8 | -17 | 6 | -5,49 | 96 | Thalamus_L | 100.0% Left_Thalamus |
| **73** | -29 | 2 | 61 | 5,95 | 96 | Frontal_Mid_2_L | 33.0% Left_Middle_Frontal_Gyrus; 18.0% Left_Superior_Frontal_Gyrus |
| **74** | 30 | -77 | -40 | -5,94 | 96 | Cerebelum_Crus2_R | 0% no_label |
| **75** | -53 | -25 | -11 | -5,57 | 96 | Temporal_Mid_L | 47.0% Left_Middle_Temporal_Gyrus_posterior_division |
| **76** | -22 | 52 | 37 | -6,23 | 96 | Frontal_Sup_2_L | 62.0% Left_Frontal_Pole |
| **77** | 2 | 52 | -8 | 5,91 | 96 | Frontal_Med_Orb_R | 51.0% Right_Frontal_Medial_Cortex; 18.0% Right_Paracingulate_Gyrus; 11.0% Right_Frontal_Pole |
| **78** | -29 | 18 | -11 | -5,46 | 96 | Insula_L | 47.0% Left_Insular_Cortex; 28.0% Left_Frontal_Orbital_Cortex |
| **79** | -39 | 49 | 1 | -6,02 | 96 | Frontal_Mid_2_L | 73.0% Left_Frontal_Pole |
| **80** | 52 | -53 | 8 | 6,32 | 96 | Temporal_Mid_R | 40.0% Right_Middle_Temporal_Gyrus_temporooccipital_part; 8.0% Right_Angular_Gyrus |
| **81** | 26 | -6 | 71 | -7,40 | 96 | Frontal_Sup_2_R | 32.0% Right_Superior_Frontal_Gyrus; 18.0% Right_Precentral_Gyrus |
| **82** | 35 | 49 | 4 | -5,77 | 82 | Frontal_Mid_2_R | 47.0% Right_Frontal_Pole |
| **83** | 26 | -44 | 13 | -6,28 | 82 | no_label | 76.0% Right_Lateral_Ventricle |

| | | | | | | | |
|---|---|---|---|---|---|---|---|
| **84** | 59 | -22 | 52 | -7,16 | 82 | Postcentral_R | 26.0% Right_Postcentral_Gyrus; 11.0% Right_Supramarginal_Gyrus_anterior_division |
| **85** | 61 | -41 | 37 | -5,53 | 82 | SupraMarginal_R | 67.0% Right_Supramarginal_Gyrus_posterior_division; 12.0% Right_Angular_Gyrus |
| **86** | -1 | -13 | 37 | -5,92 | 82 | Cingulate_Mid_L | 72.0% Left_Cingulate_Gyrus_anterior_division; 23.0% Left_Cingulate_Gyrus_posterior_division |
| **87** | -5 | 52 | 20 | -6,39 | 82 | Frontal_Sup_Medial_L | 41.0% Left_Paracingulate_Gyrus; 34.0% Left_Superior_Frontal_Gyrus; 7.0% Left_Frontal_Pole |
| **88** | -20 | -46 | -11 | -5,55 | 82 | Lingual_L | 71.0% Left_Lingual_Gyrus; 16.0% Left_Temporal_Occipital_Fusiform_Cortex |
| **89** | 11 | -84 | 35 | -5,76 | 68 | Cuneus_R | 40.0% Right_Cuneal_Cortex; 13.0% Right_Lateral_Occipital_Cortex_superior_division; 9.0% Right_Occipital_Pole |
| **90** | -1 | 30 | 13 | -5,84 | 68 | Cingulate_Ant_L | 71.0% Left_Cingulate_Gyrus_anterior_division |
| **91** | -27 | -63 | 64 | -6,88 | 68 | Parietal_Sup_L | 55.0% Left_Lateral_Occipital_Cortex_superior_division; 9.0% Left_Superior_Parietal_Lobule |
| **92** | -51 | -58 | 30 | -6,34 | 68 | Angular_L | 54.0% Left_Angular_Gyrus; 18.0% Left_Lateral_Occipital_Cortex_superior_division; 7.0% Left_Supramarginal_Gyrus_posterior_division |
| **93** | -58 | -44 | 23 | -5,57 | 68 | Temporal_Sup_L | 45.0% Left_Supramarginal_Gyrus_posterior_division; 12.0% Left_Parietal_Operculum_Cortex; 9.0% Left_Planum_Temporale; 7.0% Left_Angular_Gyrus |

***Table S4.*** *Regions whose activation changed positively or negatively when threatened with a mild electric shock. X-, Y- and Z-coordinates are in MNI space; volume is in mm.*

| # | X | Y | Z | Z-STAT | VOLUME | AAL | HARVARD-OXFORD |
|---|---|---|---|---|---|---|---|
| **1** | -8 | -87 | -1 | 28,28 | 115714 | Calcarine_L | 32.0% Left_Intracalcarine_Cortex; 22.0% Left_Lingual_Gyrus; 5.0% Left_Occipital_Pole |
| **1** | 9 | -89 | 18 | 24,70 | 115714 | Cuneus_R | 28.0% Right_Occipital_Pole; 13.0% Right_Cuneal_Cortex |
| **1** | 38 | -56 | -25 | 22,03 | 115714 | Cerebelum_6_R | 0% no_label |
| **1** | -29 | -91 | 20 | 19,88 | 115714 | Occipital_Mid_L | 44.0% Left_Occipital_Pole; 22.0% Left_Lateral_Occipital_Cortex_superior_division |
| **1** | 23 | -80 | -8 | 17,93 | 115714 | Lingual_R | 43.0% Right_Occipital_Fusiform_Gyrus; 5.0% Right_Lingual_Gyrus |
| **1** | -36 | -56 | -28 | 17,88 | 115714 | Cerebelum_6_L | 0% no_label |
| **1** | -29 | -80 | -18 | 16,53 | 115714 | Fusiform_L | 56.0% Left_Occipital_Fusiform_Gyrus; 15.0% Left_Lateral_Occipital_Cortex_inferior_division |
| **1** | 33 | -77 | 20 | 15,00 | 115714 | Occipital_Mid_R | 57.0% Right_Lateral_Occipital_Cortex_superior_division |
| **1** | 2 | -77 | -35 | 8,65 | 115714 | Cerebelum_Crus2_L | 0% no_label |
| **1** | 28 | -39 | -40 | 8,54 | 115714 | no_label | 0% no_label |
| **1** | -24 | -39 | -42 | 6,98 | 115714 | Cerebelum_10_L | 0% no_label |
| **1** | -1 | -58 | -1 | 5,86 | 115714 | Vermis_4_5 | 0% no_label |
| **2** | -3 | 4 | 61 | 20,48 | 57577 | Supp_Motor_Area_L | 81.0% Left_Juxtapositional_Lobule_Cortex_(formerly_Supplementary_Motor_Cortex); 10.0% Left_Superior_Frontal_Gyrus |
| **2** | -41 | -6 | 49 | 17,61 | 57577 | Precentral_L | 59.0% Left_Precentral_Gyrus |
| **2** | 47 | -1 | 54 | 14,83 | 57577 | Frontal_Mid_2_R | 61.0% Right_Precentral_Gyrus; 14.0% Right_Middle_Frontal_Gyrus |
| **2** | -8 | 16 | 35 | 13,64 | 57577 | Cingulate_Mid_L | 51.0% Left_Cingulate_Gyrus_anterior_division; 28.0% Left_Paracingulate_Gyrus |
| **2** | -34 | 28 | 4 | 13,19 | 57577 | Insula_L | 21.0% Left_Frontal_Operculum_Cortex; 19.0% Left_Frontal_Orbital_Cortex; 15.0% Left_Inferior_Frontal_Gyrus_pars_triangularis; 7.0% Left_Insular_Cortex |
| **2** | -60 | 4 | 4 | 12,97 | 57577 | Rolandic_Oper_L | 48.0% Left_Precentral_Gyrus; 5.0% Left_Inferior_Frontal_Gyrus_pars_opercularis |
| **2** | 26 | -3 | 66 | 10,83 | 57577 | Frontal_Sup_2_R | 40.0% Right_Superior_Frontal_Gyrus; 15.0% Right_Precentral_Gyrus; 8.0% Right_Middle_Frontal_Gyrus |
| **2** | -22 | -10 | 68 | 10,13 | 57577 | Frontal_Sup_2_L | 38.0% Left_Superior_Frontal_Gyrus; 33.0% Left_Precentral_Gyrus |
| **2** | -53 | 11 | 32 | 9,82 | 57577 | Precentral_L | 26.0% Left_Precentral_Gyrus; 23.0% Left_Inferior_Frontal_Gyrus_pars_opercularis; 17.0% Left_Middle_Frontal_Gyrus |
| **3** | 64 | -13 | 37 | -16,68 | 43955 | Postcentral_R | 65.0% Right_Postcentral_Gyrus; 8.0% Right_Supramarginal_Gyrus_anterior_division |
| **3** | 40 | -37 | 64 | -13,57 | 43955 | Postcentral_R | 51.0% Right_Postcentral_Gyrus; 20.0% Right_Superior_Parietal_Lobule |
| **3** | 38 | -8 | 13 | -12,27 | 43955 | Insula_R | 78.0% Right_Insular_Cortex; 13.0% Right_Central_Opercular_Cortex |
| **3** | 16 | -17 | 78 | -11,51 | 43955 | Precentral_R | 42.0% Right_Precentral_Gyrus; 6.0% Right_Superior_Frontal_Gyrus |
| **3** | 64 | -13 | 13 | -10,16 | 43955 | Postcentral_R | 45.0% Right_Central_Opercular_Cortex; 15.0% Right_Postcentral_Gyrus; 9.0% Right_Planum_Temporale; 6.0% Right_Planum_Polare; 5.0% Right_Parietal_Operculum_Cortex |
| **3** | -3 | -27 | 59 | -9,46 | 43955 | Paracentral_Lobule_L | 85.0% Left_Precentral_Gyrus |
| **4** | -12 | -80 | 54 | 17,83 | 19989 | Precuneus_L | 37.0% Left_Lateral_Occipital_Cortex_superior_division |
| **4** | 11 | -75 | 61 | 13,21 | 19989 | Precuneus_R | 31.0% Right_Lateral_Occipital_Cortex_superior_division |
| **4** | -5 | -58 | 66 | 13,13 | 19989 | Precuneus_L | 53.0% Left_Precuneous_Cortex; 6.0% Left_Lateral_Occipital_Cortex_superior_division; 6.0% Left_Superior_Parietal_Lobule |
| **4** | -29 | -72 | 35 | 9,46 | 19989 | Occipital_Mid_L | 73.0% Left_Lateral_Occipital_Cortex_superior_division |
| **4** | -8 | -60 | 40 | 6,89 | 19989 | Precuneus_L | 59.0% Left_Precuneous_Cortex |
| **4** | 23 | -51 | 61 | 6,63 | 19989 | Parietal_Sup_R | 45.0% Right_Superior_Parietal_Lobule |
| **5** | -65 | -13 | 32 | -12,53 | 17407 | Postcentral_L | 50.0% Left_Postcentral_Gyrus |
| **5** | -36 | -10 | 13 | -12,36 | 17407 | Insula_L | 75.0% Left_Insular_Cortex; 12.0% Left_Central_Opercular_Cortex |

| | | | | | | | |
|---|---|---|---|---|---|---|---|
| **5** | -44 | -34 | 66 | -11,81 | 17407 | Postcentral_L | 52.0% Left_Postcentral_Gyrus |
| **5** | -60 | -13 | 8 | -7,72 | 17407 | Heschl_L | 36.0% Left_Central_Opercular_Cortex; 30.0% Left_Planum_Temporale; 10.0% Left_Planum_Polare; 5.0% Left_Heschl's_Gyrus_(includes_H1_and_H2) |
| **5** | -17 | -29 | 78 | -7,45 | 17407 | Postcentral_L | 23.0% Left_Postcentral_Gyrus; 16.0% Left_Precentral_Gyrus |
| **6** | 47 | 11 | -1 | 13,77 | 8662 | Insula_R | 25.0% Right_Frontal_Operculum_Cortex; 19.0% Right_Central_Opercular_Cortex; 5.0% Right_Inferior_Frontal_Gyrus_pars_opercularis |
| **6** | 45 | 35 | -11 | 7,29 | 8662 | Frontal_Inf_Orb_2_R | 29.0% Right_Frontal_Pole; 14.0% Right_Frontal_Orbital_Cortex |
| **7** | -8 | -1 | 11 | 10,67 | 7378 | no_label | 36.0% Left_Lateral_Ventrical; 34.0% Left_Caudate; 26.0% Left_Thalamus |
| **8** | -39 | 42 | 28 | 11,22 | 5698 | Frontal_Mid_2_L | 79.0% Left_Frontal_Pole; 14.0% Left_Middle_Frontal_Gyrus |
| **9** | -55 | -41 | 32 | 17,74 | 5329 | SupraMarginal_L | 38.0% Left_Supramarginal_Gyrus_posterior_division; 28.0% Left_Supramarginal_Gyrus_anterior_division; 6.0% Left_Planum_Temporale; 6.0% Left_Parietal_Operculum_Cortex |
| **10** | -5 | -82 | 18 | -9,19 | 4823 | Cuneus_L | 29.0% Left_Cuneal_Cortex; 22.0% Left_Supracalcarine_Cortex; 12.0% Left_Intracalcarine_Cortex |
| **10** | 19 | -80 | 40 | -8,27 | 4823 | Cuneus_R | 48.0% Right_Lateral_Occipital_Cortex_superior_division; 7.0% Right_Cuneal_Cortex |
| **10** | 11 | -51 | 37 | -7,49 | 4823 | Precuneus_R | 48.0% Right_Precuneous_Cortex; 19.0% Right_Cingulate_Gyrus_posterior_division |
| **10** | 19 | -63 | 16 | -5,26 | 4823 | Calcarine_R | 35.0% Right_Supracalcarine_Cortex; 17.0% Right_Precuneous_Cortex; 15.0% Right_Cuneal_Cortex |
| **11** | 59 | -58 | 6 | -10,30 | 4727 | Temporal_Mid_R | 61.0% Right_Middle_Temporal_Gyrus_temporooccipital_part; 19.0% Right_Lateral_Occipital_Cortex_inferior_division |
| **12** | 11 | -3 | 13 | 9,42 | 4372 | no_label | 57.0% Right_Thalamus; 19.0% Right_Caudate |
| **13** | -53 | -65 | 35 | -9,74 | 2883 | Angular_L | 81.0% Left_Lateral_Occipital_Cortex_superior_division; 9.0% Left_Angular_Gyrus |
| **14** | -65 | -51 | 1 | -8,40 | 2610 | Temporal_Mid_L | 74.0% Left_Middle_Temporal_Gyrus_temporooccipital_part; 12.0% Left_Angular_Gyrus |
| **15** | 57 | -60 | 40 | -8,86 | 2541 | Parietal_Inf_R | 33.0% Right_Lateral_Occipital_Cortex_superior_division; 14.0% Right_Angular_Gyrus |
| **16** | -44 | -70 | 1 | -10,56 | 2377 | Occipital_Mid_L | 63.0% Left_Lateral_Occipital_Cortex_inferior_division |
| **17** | -22 | -8 | -16 | -7,89 | 2364 | Amygdala_L | 93.0% Left_Amygdala |
| **18** | 38 | 45 | 32 | 10,55 | 2227 | Frontal_Mid_2_R | 81.0% Right_Frontal_Pole |
| **19** | -10 | -80 | 32 | -8,56 | 1954 | Cuneus_L | 39.0% Left_Cuneal_Cortex; 17.0% Left_Precuneous_Cortex; 5.0% Left_Lateral_Occipital_Cortex_superior_division |
| **20** | -1 | -44 | 8 | 10,94 | 1858 | no_label | 13.0% Left_Cingulate_Gyrus_posterior_division |
| **21** | 28 | 16 | 54 | -8,31 | 1722 | Frontal_Mid_2_R | 45.0% Right_Middle_Frontal_Gyrus; 9.0% Right_Superior_Frontal_Gyrus |
| **22** | -36 | -46 | 37 | 12,23 | 1708 | Parietal_Inf_L | 19.0% Left_Supramarginal_Gyrus_posterior_division; 18.0% Left_Superior_Parietal_Lobule |
| **23** | 26 | -17 | -16 | -8,68 | 1325 | Hippocampus_R | 97.0% Right_Hippocampus |
| **24** | -60 | -3 | -13 | -9,36 | 1189 | Temporal_Mid_L | 45.0% Left_Middle_Temporal_Gyrus_anterior_division; 23.0% Left_Superior_Temporal_Gyrus_anterior_division |
| **25** | 35 | 9 | 28 | -11,10 | 1134 | Frontal_Inf_Oper_R | 15.0% Right_Precentral_Gyrus; 14.0% Right_Middle_Frontal_Gyrus; 12.0% Right_Inferior_Frontal_Gyrus_pars_opercularis |
| **26** | 45 | -51 | -16 | -9,03 | 1052 | Temporal_Inf_R | 48.0% Right_Temporal_Occipital_Fusiform_Cortex; 33.0% Right_Inferior_Temporal_Gyrus_temporooccipital_part |
| **27** | 16 | 56 | 28 | -7,75 | 902 | Frontal_Sup_2_R | 64.0% Right_Frontal_Pole |
| **28** | 35 | 64 | 4 | -9,07 | 847 | Frontal_Mid_2_R | 53.0% Right_Frontal_Pole |
| **29** | -27 | 28 | 44 | -7,52 | 765 | Frontal_Mid_2_L | 42.0% Left_Middle_Frontal_Gyrus; 25.0% Left_Superior_Frontal_Gyrus |
| **30** | -1 | -44 | 37 | -8,00 | 724 | Precuneus_L | 58.0% Left_Cingulate_Gyrus_posterior_division; 20.0% Left_Precuneous_Cortex |
| **31** | 64 | -37 | 30 | 7,72 | 697 | SupraMarginal_R | 59.0% Right_Supramarginal_Gyrus_posterior_division; 10.0% Right_Supramarginal_Gyrus_anterior_division; 7.0% Right_Parietal_Operculum_Cortex |
| **32** | 2 | -34 | 28 | 7,82 | 574 | no_label | 89.0% Right_Cingulate_Gyrus_posterior_division |
| **33** | 45 | 28 | 23 | -7,94 | 533 | Frontal_Mid_2_R | 43.0% Right_Middle_Frontal_Gyrus; 16.0% Right_Inferior_Frontal_Gyrus_pars_triangularis |
| **34** | 2 | 40 | -11 | -7,78 | 506 | Frontal_Med_Orb_R | 51.0% Right_Paracingulate_Gyrus; 37.0% Right_Frontal_Medial_Cortex; 6.0% Right_Cingulate_Gyrus_anterior_division |

| | | | | | | | |
|---|---|---|---|---|---|---|---|
| **35** | 23 | -25 | -6 | 7,96 | 451 | no_label | 5.0% Right_Thalamus |
| **36** | -22 | -27 | -6 | 7,72 | 437 | Hippocampus_L | 5.0% Left_Hippocampus |
| **37** | -8 | -48 | 1 | 9,99 | 369 | Lingual_L | 41.0% Left_Cingulate_Gyrus_posterior_division; 17.0% Left_Lingual_Gyrus |
| **38** | 16 | 33 | 59 | -7,51 | 355 | Frontal_Sup_2_R | 31.0% Right_Superior_Frontal_Gyrus; 7.0% Right_Frontal_Pole |
| **39** | 11 | -53 | 49 | 6,92 | 342 | Precuneus_R | 43.0% Right_Precuneous_Cortex; 5.0% Right_Superior_Parietal_Lobule |
| **40** | -24 | -99 | -6 | -9,40 | 328 | Occipital_Inf_L | 68.0% Left_Occipital_Pole |
| **41** | 2 | 14 | -18 | -6,95 | 328 | no_label | 71.0% Right_Subcallosal_Cortex |
| **42** | 11 | -34 | 4 | -6,45 | 314 | no_label | 70.0% Right_Thalamus; 9.0% Right_Hippocampus |
| **43** | -44 | -56 | -13 | -7,59 | 314 | Fusiform_L | 39.0% Left_Inferior_Temporal_Gyrus_temporooccipital_part; 31.0% Left_Temporal_Occipital_Fusiform_Cortex; 6.0% Left_Lateral_Occipital_Cortex_inferior_division |
| **44** | 19 | -1 | -18 | -7,12 | 287 | Amygdala_R | 88.0% Right_Amygdala; 7.0% Right_Parahippocampal_Gyrus_anterior_division |
| **45** | -3 | 23 | -20 | -6,26 | 273 | Rectus_L | 82.0% Left_Subcallosal_Cortex |
| **46** | -44 | -3 | -6 | -6,55 | 273 | Insula_L | 21.0% Left_Insular_Cortex; 16.0% Left_Planum_Polare |
| **47** | -3 | 52 | 13 | -6,44 | 246 | Frontal_Sup_Medial_L | 72.0% Left_Paracingulate_Gyrus; 11.0% Left_Superior_Frontal_Gyrus; 7.0% Left_Frontal_Pole |
| **48** | 2 | -53 | -35 | 7,53 | 246 | Vermis_9 | 0% no_label |
| **49** | 50 | -25 | -1 | 7,49 | 219 | Temporal_Sup_R | 34.0% Right_Superior_Temporal_Gyrus_posterior_division; 9.0% Right_Middle_Temporal_Gyrus_posterior_division |
| **50** | -51 | 6 | -28 | -6,07 | 219 | Temporal_Mid_L | 68.0% Left_Temporal_Pole |
| **51** | 69 | -13 | -13 | -6,70 | 219 | Temporal_Mid_R | 54.0% Right_Middle_Temporal_Gyrus_posterior_division; 5.0% Right_Middle_Temporal_Gyrus_anterior_division |
| **52** | -44 | 16 | -35 | -6,23 | 205 | Temporal_Pole_Mid_L | 75.0% Left_Temporal_Pole |
| **53** | 54 | -10 | -30 | -6,44 | 205 | Temporal_Inf_R | 25.0% Right_Middle_Temporal_Gyrus_posterior_division; 21.0% Right_Inferior_Temporal_Gyrus_posterior_division; 15.0% Right_Inferior_Temporal_Gyrus_anterior_division; 8.0% Right_Middle_Temporal_Gyrus_anterior_division |
| **54** | -67 | -29 | -18 | -7,19 | 205 | Temporal_Inf_L | 57.0% Left_Middle_Temporal_Gyrus_posterior_division; 15.0% Left_Inferior_Temporal_Gyrus_posterior_division |
| **55** | -10 | 52 | 47 | -6,12 | 205 | no_label | 20.0% Left_Frontal_Pole |
| **56** | -17 | -63 | -4 | -6,55 | 191 | Lingual_L | 37.0% Left_Lingual_Gyrus; 5.0% Left_Temporal_Occipital_Fusiform_Cortex |
| **57** | -29 | -20 | 1 | 6,34 | 178 | no_label | 82.0% Left_Putamen |
| **58** | 59 | -17 | -8 | -6,40 | 164 | Temporal_Mid_R | 45.0% Right_Middle_Temporal_Gyrus_posterior_division; 17.0% Right_Superior_Temporal_Gyrus_posterior_division |
| **59** | -10 | -25 | 40 | 6,58 | 164 | Cingulate_Mid_L | 67.0% Left_Cingulate_Gyrus_posterior_division; 12.0% Left_Precentral_Gyrus |
| **60** | -5 | 16 | -13 | -6,31 | 150 | Olfactory_L | 84.0% Left_Subcallosal_Cortex |
| **61** | -34 | 37 | -13 | 6,47 | 150 | OFCant_L | 52.0% Left_Frontal_Pole; 42.0% Left_Frontal_Orbital_Cortex |
| **62** | 47 | 14 | 49 | -6,67 | 150 | Frontal_Mid_2_R | 65.0% Right_Middle_Frontal_Gyrus |
| **63** | 54 | 9 | -30 | -6,58 | 150 | Temporal_Pole_Mid_R | 66.0% Right_Temporal_Pole |
| **64** | -51 | -48 | 18 | -6,55 | 150 | Temporal_Mid_L | 23.0% Left_Supramarginal_Gyrus_posterior_division; 20.0% Left_Angular_Gyrus |
| **65** | 45 | -39 | 6 | 6,63 | 150 | Temporal_Sup_R | 20.0% Right_Supramarginal_Gyrus_posterior_division; 16.0% Right_Middle_Temporal_Gyrus_temporooccipital_part; 9.0% Right_Middle_Temporal_Gyrus_posterior_division |
| **66** | 26 | 59 | 18 | -6,38 | 137 | Frontal_Sup_2_R | 84.0% Right_Frontal_Pole |
| **67** | -34 | -65 | 59 | 7,42 | 137 | Parietal_Sup_L | 55.0% Left_Lateral_Occipital_Cortex_superior_division |
| **68** | -24 | 33 | -13 | 6,01 | 137 | OFCpost_L | 53.0% Left_Frontal_Orbital_Cortex; 22.0% Left_Frontal_Pole |
| **69** | -10 | -53 | 35 | -6,62 | 137 | Precuneus_L | 47.0% Left_Precuneous_Cortex; 17.0% Left_Cingulate_Gyrus_posterior_division |
| **70** | -24 | -39 | 1 | -6,11 | 123 | Hippocampus_L | 67.0% Left_Hippocampus; 7.0% Left_Lateral_Ventrical |
| **71** | 14 | -25 | 40 | 6,34 | 123 | Cingulate_Mid_R | 48.0% Right_Cingulate_Gyrus_posterior_division; 26.0% Right_Precentral_Gyrus |
| **72** | 11 | -13 | -4 | 5,94 | 123 | no_label | 9.0% Right_Thalamus |
| **73** | 50 | -1 | -4 | -6,19 | 123 | no_label | 55.0% Right_Planum_Polare; 7.0% Right_Heschl's_Gyrus_(includes_H1_and_H2) |

| | | | | | | | |
|---|---|---|---|---|---|---|---|
| **74** | 2 | 28 | 13 | -6,96 | 123 | no_label | 51.0% Right_Cingulate_Gyrus_anterior_division; 5.0% Left_Cingulate_Gyrus_anterior_division |
| **75** | -10 | -48 | 68 | -6,38 | 109 | Precuneus_L | 36.0% Left_Postcentral_Gyrus; 18.0% Left_Superior_Parietal_Lobule; 14.0% Left_Precuneous_Cortex |
| **76** | -27 | -25 | -30 | 5,87 | 109 | Fusiform_L | 24.0% Left_Parahippocampal_Gyrus_posterior_division; 19.0% Left_Parahippocampal_Gyrus_anterior_division; 17.0% Left_Temporal_Fusiform_Cortex_posterior_division |
| **77** | 30 | 14 | -35 | -5,74 | 109 | Temporal_Pole_Mid_R | 71.0% Right_Temporal_Pole |
| **78** | 16 | -58 | -6 | -6,07 | 109 | Lingual_R | 74.0% Right_Lingual_Gyrus |
| **79** | 2 | -39 | -40 | 5,54 | 109 | no_label | 97.0% Brain-Stem |
| **80** | 4 | 25 | 66 | 6,00 | 109 | no_label | 9.0% Right_Superior_Frontal_Gyrus |
| **81** | 45 | -15 | 1 | -6,43 | 96 | Temporal_Sup_R | 36.0% Right_Heschl's_Gyrus_(includes_H1_and_H2); 23.0% Right_Planum_Polare |
| **82** | 2 | 59 | -16 | -7,11 | 82 | Rectus_R | 73.0% Right_Frontal_Pole |
| **83** | 35 | 18 | -37 | -6,93 | 82 | Temporal_Pole_Mid_R | 55.0% Right_Temporal_Pole |
| **84** | -34 | 16 | 59 | -5,64 | 82 | Frontal_Mid_2_L | 63.0% Left_Middle_Frontal_Gyrus |
| **85** | -22 | -68 | 18 | -5,82 | 82 | no_label | 16.0% Left_Cuneal_Cortex; 12.0% Left_Supracalcarine_Cortex; 6.0% Left_Precuneous_Cortex |
| **86** | -17 | -25 | -1 | -6,26 | 82 | Thalamus_L | 77.0% Left_Thalamus |
| **87** | -3 | 40 | 8 | -5,88 | 82 | Cingulate_Ant_L | 91.0% Left_Cingulate_Gyrus_anterior_division; 8.0% Left_Paracingulate_Gyrus |
| **88** | -48 | 25 | 32 | 6,16 | 68 | Frontal_Mid_2_L | 66.0% Left_Middle_Frontal_Gyrus; 5.0% Left_Inferior_Frontal_Gyrus_pars_triangularis |
| **89** | 19 | -89 | 42 | 5,96 | 68 | no_label | 21.0% Right_Occipital_Pole; 14.0% Right_Lateral_Occipital_Cortex_superior_division |
| **90** | 9 | -46 | 1 | 7,08 | 68 | Vermis_4_5 | 39.0% Right_Cingulate_Gyrus_posterior_division; 26.0% Right_Lingual_Gyrus |
| **91** | -65 | -56 | 23 | -6,45 | 68 | no_label | 6.0% Left_Angular_Gyrus |
| **92** | 45 | 14 | -28 | -5,50 | 68 | Temporal_Pole_Sup_R | 62.0% Right_Temporal_Pole |
| **93** | -39 | 33 | 44 | 6,19 | 68 | Frontal_Mid_2_L | 29.0% Left_Middle_Frontal_Gyrus |
| **94** | 69 | -27 | 28 | 5,57 | 68 | SupraMarginal_R | 17.0% Right_Supramarginal_Gyrus_anterior_division |
| **95** | -1 | -37 | 76 | 6,28 | 68 | no_label | 0% no_label |

*Table S5. Regions whose activation changed positively or negatively when threatened with a mild electric shock, compared to not being threatened, during regulation towards SN. X-, Y- and Z-coordinates are in MNI space; volume is in mm.*

| # | X | Y | Z | Z-STAT | VOLUME | AAL | HARVARD-OXFORD |
|---|---|---|---|---|---|---|---|
| **1** | 11 | -82 | 4 | 21,25 | 46633 | Calcarine_R | 47.0% Right_Intracalcarine_Cortex; 14.0% Right_Lingual_Gyrus |
| **1** | -12 | -77 | -8 | 16,78 | 46633 | Lingual_L | 57.0% Left_Lingual_Gyrus; 9.0% Left_Occipital_Fusiform_Gyrus |
| **1** | -10 | -96 | 16 | 14,72 | 46633 | Cuneus_L | 56.0% Left_Occipital_Pole |
| **1** | 16 | -94 | 28 | 11,71 | 46633 | Occipital_Sup_R | 47.0% Right_Occipital_Pole; 7.0% Right_Lateral_Occipital_Cortex_superior_division |
| **1** | 23 | -68 | -13 | 10,42 | 46633 | Fusiform_R | 54.0% Right_Occipital_Fusiform_Gyrus; 23.0% Right_Lingual_Gyrus |
| **1** | -34 | -84 | 6 | 8,33 | 46633 | Occipital_Mid_L | 41.0% Left_Lateral_Occipital_Cortex_inferior_division; 23.0% Left_Lateral_Occipital_Cortex_superior_division |
| **1** | 33 | -75 | 20 | 8,16 | 46633 | Occipital_Mid_R | 53.0% Right_Lateral_Occipital_Cortex_superior_division |
| **1** | -29 | -53 | -8 | 8,04 | 46633 | Fusiform_L | 55.0% Left_Temporal_Occipital_Fusiform_Cortex; 20.0% Left_Lingual_Gyrus |
| **1** | 40 | -87 | -4 | 6,24 | 46633 | Occipital_Inf_R | 52.0% Right_Lateral_Occipital_Cortex_inferior_division; 15.0% Right_Occipital_Pole |
| **2** | 2 | -25 | 28 | 8,54 | 2145 | no_label | 69.0% Right_Cingulate_Gyrus_posterior_division |
| **3** | 7 | -77 | 54 | 7,13 | 1558 | Precuneus_R | 30.0% Right_Precuneous_Cortex; 14.0% Right_Lateral_Occipital_Cortex_superior_division |
| **3** | 16 | -60 | 32 | 6,75 | 1558 | Precuneus_R | 44.0% Right_Precuneous_Cortex |
| **4** | 35 | 25 | 6 | 7,68 | 1025 | Insula_R | 48.0% Right_Frontal_Operculum_Cortex; 14.0% Right_Insular_Cortex; 8.0% Right_Frontal_Orbital_Cortex; 6.0% Right_Inferior_Frontal_Gyrus_pars_triangularis |
| **5** | -12 | -68 | 37 | 7,48 | 915 | Occipital_Sup_L | 24.0% Left_Precuneous_Cortex; 5.0% Left_Cuneal_Cortex |
| **6** | 50 | -44 | 42 | 7,16 | 874 | SupraMarginal_R | 41.0% Right_Supramarginal_Gyrus_posterior_division; 25.0% Right_Angular_Gyrus |
| **7** | 11 | -44 | 35 | 6,36 | 314 | Cingulate_Mid_R | 47.0% Right_Cingulate_Gyrus_posterior_division; 25.0% Right_Precuneous_Cortex |
| **8** | 2 | 25 | 56 | 6,62 | 246 | Frontal_Sup_Medial_L | 42.0% Right_Superior_Frontal_Gyrus; 6.0% Left_Superior_Frontal_Gyrus |
| **9** | -1 | -51 | -35 | 6,70 | 246 | Vermis_10 | 0% no_label |
| **10** | 33 | 59 | 16 | 6,02 | 178 | Frontal_Sup_2_R | 90.0% Right_Frontal_Pole |
| **11** | -34 | 30 | 1 | 6,20 | 137 | Frontal_Inf_Tri_L | 34.0% Left_Frontal_Orbital_Cortex; 25.0% Left_Inferior_Frontal_Gyrus_pars_triangularis |
| **12** | -58 | -1 | 6 | 6,80 | 137 | Rolandic_Oper_L | 36.0% Left_Central_Opercular_Cortex; 32.0% Left_Precentral_Gyrus; 13.0% Left_Planum_Polare |
| **13** | 9 | -53 | 49 | 5,88 | 109 | Precuneus_R | 61.0% Right_Precuneous_Cortex |
| **14** | 47 | -53 | 37 | 5,84 | 109 | Angular_R | 49.0% Right_Angular_Gyrus |
| **15** | -1 | -39 | 1 | 5,75 | 82 | no_label | 0% no_label |
| **16** | 14 | -68 | 49 | 5,35 | 82 | Parietal_Sup_R | 29.0% Right_Lateral_Occipital_Cortex_superior_division; 26.0% Right_Precuneous_Cortex |
| **17** | 57 | -41 | 30 | 5,83 | 82 | SupraMarginal_R | 52.0% Right_Supramarginal_Gyrus_posterior_division; 9.0% Right_Angular_Gyrus |
| **18** | 4 | -63 | 42 | 5,65 | 68 | Precuneus_R | 92.0% Right_Precuneous_Cortex |
| **19** | 30 | 21 | -11 | 6,18 | 68 | Insula_R | 63.0% Right_Frontal_Orbital_Cortex; 23.0% Right_Insular_Cortex |
| **20** | 42 | -65 | 52 | 5,54 | 68 | Angular_R | 70.0% Right_Lateral_Occipital_Cortex_superior_division |

*Table S6. Regions whose activation changed positively or negatively when threatened with a mild electric shock, compared to not being threatened, during regulation towards ECN. X-, Y- and Z-coordinates are in MNI space; volume is in mm.*

| # | X | Y | Z | Z-STAT | VOLUME | AAL | HARVARD-OXFORD |
|---|---|---|---|---|---|---|---|
| **1** | -8 | -87 | -1 | 20,51 | 36453 | Calcarine_L | 32.0% Left_Intracalcarine_Cortex; 22.0% Left_Lingual_Gyrus; 5.0% Left_Occipital_Pole |
| **1** | 9 | -89 | 18 | 18,81 | 36453 | Cuneus_R | 28.0% Right_Occipital_Pole; 13.0% Right_Cuneal_Cortex |
| **1** | 23 | -68 | -4 | 9,91 | 36453 | Lingual_R | 31.0% Right_Lingual_Gyrus; 27.0% Right_Occipital_Fusiform_Gyrus |
| **1** | -29 | -91 | 20 | 8,78 | 36453 | Occipital_Mid_L | 44.0% Left_Occipital_Pole; 22.0% Left_Lateral_Occipital_Cortex_superior_division |
| **1** | -29 | -77 | -16 | 7,95 | 36453 | Fusiform_L | 72.0% Left_Occipital_Fusiform_Gyrus; 9.0% Left_Lateral_Occipital_Cortex_inferior_division |
| **1** | -29 | -53 | -8 | 7,11 | 36453 | Fusiform_L | 55.0% Left_Temporal_Occipital_Fusiform_Cortex; 20.0% Left_Lingual_Gyrus |
| **1** | 33 | -77 | 20 | 7,03 | 36453 | Occipital_Mid_R | 57.0% Right_Lateral_Occipital_Cortex_superior_division |
| **2** | 30 | 21 | -11 | 6,56 | 888 | Insula_R | 63.0% Right_Frontal_Orbital_Cortex; 23.0% Right_Insular_Cortex |
| **3** | 38 | -27 | 68 | -6,78 | 874 | Precentral_R | 45.0% Right_Postcentral_Gyrus; 20.0% Right_Precentral_Gyrus |
| **4** | 66 | -13 | 35 | -6,05 | 342 | Postcentral_R | 40.0% Right_Postcentral_Gyrus |
| **5** | 4 | -20 | 28 | 6,28 | 287 | no_label | 51.0% Right_Cingulate_Gyrus_posterior_division; 10.0% Right_Cingulate_Gyrus_anterior_division |
| **6** | 54 | -41 | 40 | 6,16 | 260 | SupraMarginal_R | 50.0% Right_Supramarginal_Gyrus_posterior_division; 10.0% Right_Angular_Gyrus |
| **7** | -1 | -29 | 59 | -5,86 | 191 | no_label | 38.0% Left_Precentral_Gyrus |
| **8** | 30 | -77 | -6 | 6,61 | 191 | Fusiform_R | 38.0% Right_Occipital_Fusiform_Gyrus; 8.0% Right_Lateral_Occipital_Cortex_inferior_division |
| **9** | 50 | -39 | 11 | -5,82 | 164 | Temporal_Sup_R | 29.0% Right_Supramarginal_Gyrus_posterior_division; 14.0% Right_Middle_Temporal_Gyrus_temporooccipital_part |
| **10** | 14 | -20 | 76 | -6,06 | 150 | Precentral_R | 63.0% Right_Precentral_Gyrus |
| **11** | 23 | -17 | -16 | -6,28 | 123 | Hippocampus_R | 93.0% Right_Hippocampus |
| **12** | -34 | -82 | 6 | 5,68 | 109 | Occipital_Mid_L | 35.0% Left_Lateral_Occipital_Cortex_inferior_division; 16.0% Left_Lateral_Occipital_Cortex_superior_division |
| **13** | 26 | -37 | 71 | -5,50 | 96 | Postcentral_R | 52.0% Right_Postcentral_Gyrus; 11.0% Right_Superior_Parietal_Lobule |
| **14** | -58 | 2 | 4 | 6,67 | 96 | Rolandic_Oper_L | 33.0% Left_Precentral_Gyrus; 28.0% Left_Central_Opercular_Cortex; 13.0% Left_Planum_Polare |
| **15** | 47 | -20 | 61 | -6,06 | 68 | Postcentral_R | 48.0% Right_Postcentral_Gyrus; 12.0% Right_Precentral_Gyrus |
| **16** | 21 | -37 | 78 | -5,64 | 68 | Postcentral_R | 19.0% Right_Postcentral_Gyrus |
